# Magnetically controlled tension of cytoskeletal elements with magnetic nanoparticles affects the expression of signaling pathway genes associated with cytoskeletal elements

**DOI:** 10.1101/2025.05.08.652598

**Authors:** Varvara Pozdina, Olga Karavashkova, Artem Minin, Alexandra Maltseva, Ilya Zubarev

## Abstract

Interactions with the extracellular environment and biological responses are based on biochemical pathways. Cytoskeletal reorganization is a dynamic process accompanied by filament polymerization and depolymerization, that allows the cell to effectively perceive and respond to external mechanical stimuli by altering their biomechanical properties. The contribution of mechanical deformations of individual cytoskeletal elements to changes in intracellular signaling has not been covered in the existing literature. In our article, we investigated changes in gene expression after regulated cytoskeletal deformation using a constant magnetic field and magnetic nanoparticles associated with antibodies to cytoskeletal proteins (vimentin, beta-actin and acetylated tubulin). For the first time, we have identified and described the biological pathways involved in the regulation of mechanical deformations of cytoskeletal elements.

## 1. Introduction

Human cells sense the extracellular matrix and surrounding cells through a dynamic system of cell-matrix and cell-cell contacts. Information about the rigidity/hardness of the environment, chemical composition, biochemical properties and interaction strength is encoded through the membrane connection with the environment. The cell’s ability to stretch, contract and deform locally is integral to determining its physiological responses. Information about the cellular environment must be encoded into intracellular biochemical signaling pathways. There are many signaling pathways in cells that are responsible for ligand recognition and mechanosensitivity. The process of cell recognition and the transformation of mechanical stimuli into a biochemical response involves signalling pathways such as FAK-ERK1/2 and Rho/ROCK signalling pathways. These activate cell contractility, remodelling of the extracellular matrix and cytoskeleton, or activate the YAP/TAZ signalling cascade and downstream transcription factors [1, 2].

It has been demonstrated that nanoparticles have the capacity to regulate the differentiation and physiology of stem cells. The intensity of the magnetic field has been shown to regulate stem cell differentiation along the osteogenic or adipogenic pathway [3]. This approach has even been extended to the mechanical regulation of neuronal cell function [4] and the stimulation of stem cell differentiation into neuronal lineages [5]. Furthermore, magnetic nanoparticle-driven activation promotes asymmetric tissue growth and proliferation, enhancing patterning in human nerve cells [6].

One simple method of altering the cytoskeleton is to subject the cells to compression and stretching during in vitro cultivation. If the membrane and cytoplasm are deformed, this will also result in the deformation of the cytoskeleton components, i.e. microtubules, actin microfilaments and intermediate filaments. Such mechanical deformation can lead to activation of intracellular signalling pathways and physiological effects in cells.

At the same time, the role of individual filaments in perceiving the cellular environment and participating in intracellular signaling is mainly based on the use of fibril polymerization inhibitors or introducing mutations that change the structure of proteins. These methods of cytoskeleton fibril modifications are effective in determining the role of pathogenic and opportunistic mutations in human health. However, they are not able to determine the contribution of individual cytoskeletal elements to the mechanosensitivity of the cell. The ability to mechanically deform a specific component of the cytoskeleton without altering the overall shape of the cell, along with the capability to identify the intracellular signaling pathways associated with each component, has been a long-standing research goal. In our study, we leverage magnetic nanoparticles to assess the impact of mechanical forces on a distinct component of the cytoskeleton, exploring its influence on intracellular signaling pathways.

This information is essential for understanding how a human cell determines its environment and what mechanisms underlie the programming of the cellular response afterwards.

## 2. Materials and Methods

### 2.1 Nanoparticle synthesis

In our experiments, we used magnetic Fe3O4 nanoparticles. The surface of nanoparticles contains carboxyl groups, which provide hydrophilic properties and the potential for cross-linking with proteins.

The synthesis of nanoparticles was performed according to the methods described in the articles [7] and [8]. A saturated aqueous solution of ammonia was added to aqueous solutions of FeSO4×7H2O and FeCl3×6H2O while stirring and sonicating in an ultrasonic bath at a temperature of 40°C. After 10 minutes, the nanoparticles were precipitated using a magnet, washed with water, and dispersed in water. To modify the surface of the nanoparticles, a PMIDA solution was added to them. The colloidal solution was prepared by first mixing it on an automatic stirrer. Then, it was centrifuged at 25000 rpm for 15 minutes. Following this, the solution was washed again with water and then dispersed in water. The nanoparticle suspension was filtered through a syringe filter with a pore diameter of 0.22 μm (JetBiofill, China) and diluted with culture medium.

### 2.2 Nanoparticle modification

The nanoparticles were modified with protein G, which allows them to conjugate with a heavy chain of an antibody. The conjugation of nanoparticles with protein G was performed via carbodiimide reaction [9]. The modification was carried out in two steps. First, the carboxyl groups of the nanoparticles were activated with a solution of EDC (1-Ethyl-3-(3-dimethylaminopropyl)carbodiimide) and sulfo-NHS (N-hydroxysulfosuccinimide) in 0.1M MES (2-(N-morpholino)ethanesulfonic acid) buffer (pH 5.0). A total of 1.8 mg of nanoparticles were dissolved in 45 µL of distilled water. Then, 120 microliters of the EDC/sulfo-NHS solution in MES (3 milligrams of EDC and 6 milligrams of sulfo-NHS) were added to the solution. The solution was then treated with an ultrasonic disperser and incubated for 15 minutes at room temperature. To remove unbound reagents, the solution was centrifuged for 5 minutes (15000g), the supernatant was discarded, and 450 µg of protein G dissolved in 600 µL of borate buffer (0.4 M H3BO3, 70 mM Na2B4O7·10H2O, pH 8.0) was added to the pellet. The nanoparticles were incubated overnight at 4°C, then centrifuged to remove unbound protein G and dispersed in water. The final suspension concentration was 17.25 mg/mL.

### 2.3 Cross-linking nanoparticles with antibodies

Beta-actin antibody (ThermoFisher, PA1-183), acetylated alpha-tubulin antibody (Sigma-Aldrich, T7451) and vimentin antibody were used.

Nanoparticle solution was pre-sonicated with an ultrasound bath and then mixed with antibodies and PBS (10 μl of protein G-modified nanoparticle solution, 4 μl of antibodies and 190 μl of sterile PBS), incubated for 15-30 minutes at RT, centrifuged for 5 minutes (15000 rpm) to remove unbound proteins and dissolved in 200 μl PBS. The resulting solution was diluted to 1 ml with culture medium without FBS. The microtube containing the final nanoparticle solution was sonicated prior to addition to cells to remove any nanoparticle aggregates.

### 2.4 Cell culture

We used human bone marrow mesenchymal stem cells (174H) obtained from a patient. The studies were conducted in accordance with the Declaration of Helsinki and with the approval of the local ethics committee. The cells were cultured in DMEM culture medium (PANECO, C410p) supplemented with 10% FBS (Biosera), L-glutamine, and antibiotics: penicillin at 100 U/ml and streptomycin at 100 μg/ml (Gibco). The cells were incubated in an incubator at 37oC with a CO2 concentration of 5% in T25 flasks (Jet Biofil, China) and in confocal Petri dishes with a diameter of 35 mm (Jet Biofil, China). For cell passaging, a 0.25% solution of trypsin-EDTA (PANECO, R036p) and Versene solution (PANECO, R080p) were used.

### 2.5 Experimental design

Cells were pretreated for 60 min with 1 ml of 100mM monodansylcadaverine solution in FBS-free medium to inhibit endocytosis [10]. Then, the cells were incubated with antibody-modified nanoparticle solution on a vertical magnetic system for 60 min and then without a magnetic system for 30 min [11]. After incubation with nanoparticles, cells were washed with PBS 1-2 times to remove nanoparticles not taken up by cells, then removed from the Petri dish using Versene and trypsin solution, transferred to a new confocal Petri dish and left in the incubator for 1-1.5 hours to allow cells to attach to the bottom of the culture dish. Immediately after the cells adhered, the cultural dishes with cells were placed on a lateral magnetic system and incubated for 20 hours.

The following samples were used in the work: cells incubated with nanoparticles modified with antibodies to beta-actin (nanoparcticles_actin), to vimentin (nanoparcticles_vim2) and to acetylated alpha-tubulin (nanoparcticles_tubulin), as well as cells incubated with nanoparticles that were not subjected to modification (nanoparticles), and cells to which nanoparticles were not added (control).

### 2.6 RNA isolation

After incubation on the lateral magnetic system, the cells were washed with PBS, detached from the Petri dish using Versene and trypsin solutions, and centrifuged to separate them from the culture medium. The cell pellets were washed again with PBS, the liquid was removed, and the tubes were frozen in liquid nitrogen. Samples were stored at −80 °C.

Total RNA was isolated using the HiPure Total RNA Kit (Magen) and DNase I Set (Magen). RNA concentration and quality were assessed using the Equalbit RNA BR Assay Kit (Vazyme) on a Qubit 4 fluorometer and the RNA ScreenTape assay on the Tapestation 4150 system (Agilent), respectively.

### 2.7 RNA sequencing

RNA sequencing was performed as described in [12]. Library preparation and ribosomal RNA depletion were carried out using the KAPA RNA HyperPrep Kit with rRNA Erase (HMR only). Library concentrations were measured using the Equalbit 1× dsDNA HS Assay Kit (Vazyme), and quality was assessed with the Agilent Tapestation (Agilent).

RNA sequencing was conducted using the NovaSeq 6000 S4 Reagent Kit v1.5 (200 cycles) on the Illumina NovaSeq 6000 system in a paired-end mode with a read length of 2×100 bp, generating at least 30 million raw reads per sample. Data quality was monitored using Illumina SAV, and demultiplexing was performed with Illumina Bcl2fastq2 v2.17 software.

### 2.8 RNA-Seq Data Processing

FASTQ files from RNA sequencing were processed using the STAR aligner [13] in “GeneCounts” mode with the Ensembl human transcriptome annotation (GRCh38 genome build, GRCh38.89 transcript annotation). Ensembl gene IDs were converted to HGNC gene symbols using the Complete HGNC dataset [dataset] [14].

A total of 36,596 genes with HGNC identifiers were quantified. VST normalization of the RNA-seq data was performed using DESeq2 [15], with a pseudocount of 1 added to the normalized counts.

Principal component analysis (PCA) was performed using TPM gene expression values with the ggplot2 software (v. 3.5.2).

### 2.9 Differential expression analysis

Differential expression analysis was performed using DESeq2 (v. 1.47.5). Each sample group: nanoparticles with actin (nanoparticles_actin), nanoparticles with tubulin (nanoparticles_tubulin), nanoparticles with vim2 (nanoparticles_vim2), and empty nanoparticles (nanoparticles) was compared against untreated control cells (control), and differentially expressed genes (DEGs) were identified.

DEGs were determined using the thresholds: −log10 (p value) > 30 and |log2 (Fold Change)| > 1.

Identified DEGs from each comparison were used to generate heatmaps, Volcano plots, and UpSet plots to analyze the shared and unique sets of DEGs among the groups.

### 2.10 Analysis of pathway activity and enrichment using GO (Gene Ontology), KEGG (Kyoto Encyclopedia of Genes and Genomes), and FGSEA

For functional annotation of DEGs, Gene Ontology (GO) analysis was performed using the clusterProfiler package (v. 4.8.2) for DEGs passing the thresholds: −log10 (p value) > 30 and |log2 (Fold Change)| > 1. Biological Process categories were used to analyze activated and suppressed processes.

To identify disrupted molecular pathways due to nanoparticle exposure, KEGG pathway enrichment analysis was conducted using the R package clusterProfiler (v. 4.8.2). For each experimental group (nanoparticles, nanoparticles_tubulin, nanoparticles_vim2, nanoparticles_actin), DEGs passing the filtering thresholds were analyzed to determine activated and suppressed pathways. Network graphs of gene interactions were also constructed.

To further explore the functional significance of the differentially expressed genes (DEGs), gene set enrichment analysis (GSEA) was performed using the fgsea package (v. 1.22.0) in R. This method enables the detection of changes in predefined gene sets, such as those related to biological processes, molecular functions, or pathways, providing insights into potential biological consequences of gene expression changes. The analysis utilized a ranked list of genes (VST-normalized) sorted by differential expression statistics (log fold change and p-value) obtained from the DESeq2 analysis. The analysis was performed separately for each experimental group using the Hallmark database for pathway enrichment.

## 3. Results

The cell line used in this study has a mesenchymal origin, and therefore the expected changes in biochemical and intracellular signaling pathways are limited by the phenotypic characteristics of the cells.

### 3.1 Characterization of the Transcriptomic Data

Transcriptomic data analysis from RNA sequencing was performed to assess global changes in gene expression. FASTQ files were processed using the STAR aligner in “GeneCounts” mode with annotation based on the human transcriptome from Ensembl (GRCh38 genome build, GRCh38.89 annotation). Ensembl gene identifiers were converted to official HGNC symbols using the Complete HGNC database [14] (version dated July 13, 2017), enabling the identification of expression levels for 58,233 genes.

The DESeq2 package was used to normalize the read counts, then a pseudocount of 1 was added to the normalized values. Principal component analysis (PCA) was performed based on the TPM values and sample-to-sample distance heatmap analysis to visualize the global structure of the data and assess sample reproducibility(Figure 1).

**Figure 1.**
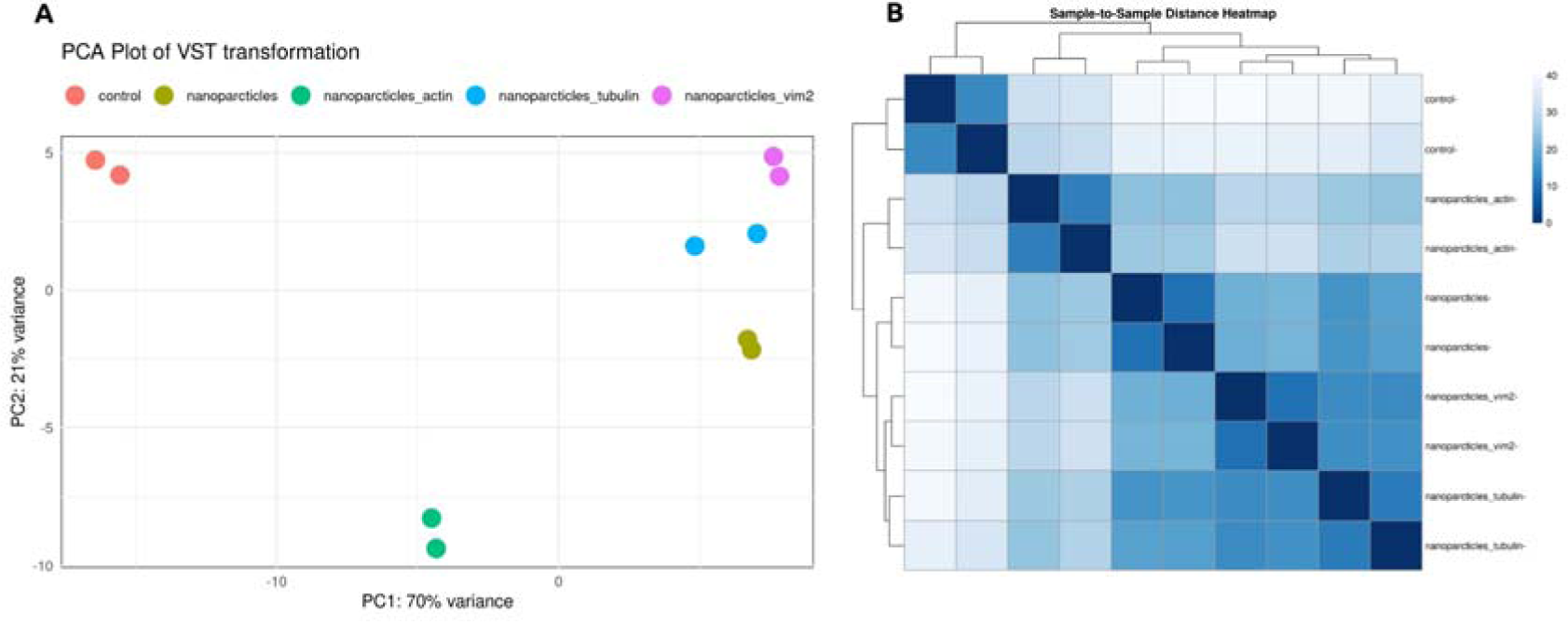
Principal component analysis (PCA) plot and Sample-to-Sample Distance Heatmap plot showing the clustering of VST (variance stabilizing transformation) transcriptomic data: A - Dots are colored by groups; B - Clustering of 10 RNA-seq samples.

PCA analysis revealed a clear separation of samples according to treatment conditions (see Fig. 1.A). The first principal component (PC1), accounting for 70% of the data variance, distinctly separates the group of intact cells (control) from cells exposed to nanoparticles. The second component (PC2), accounting for 21% of the variance, further differentiates the nanoparticle subgroups based on the type of exposure (nanoparticles, nanoparticles_tubulin, and nanoparticles_vim2). These results suggest significant changes in gene expression in experimental groups, indicating a specific molecular response of the cells to the nanoparticles.

The sample-to-sample distance heatmap was calculated using all normalized gene expression values (Figure 1.B). Distances were computed based on the Euclidean metric after variance-stabilizing transformation (VST), which reduced the influence of highly expressed genes and minimized variance associated with technical effects.

The resulting heatmap demonstrates clear clustering of samples according to experimental groups, confirming high reproducibility and distinct transcriptional differences. The closest distances are observed between replicates of the same type, whereas samples exposed to different treatments form separate clusters.

### 3.2 Differential gene expression analysis

Differential gene expression analysis was performed using the DESeq2 package (v. 1.47.5). The four experimental groups (nanoparticles_actin, nanoparticles_tubulin, nanoparticles_vim2, nanoparticles) were compared against untreated cells (control). Differentially expressed genes (DEGs) were identified using the thresholds of −log10(p-value) > 30 and |log2(Fold Change)| > 1.

The total number of DEGs in each comparison group was: 384 in the nanoparticles group, 306 in the nanoparticles_tubulin group, 405 in the nanoparticles_vim2 group, and 181 in the nanoparticles_actin group (see Table 1). The complete list of filtered genes is provided in <u>Supplementary 1</u>.

**Table 1.** Number of DEGs passing the established filtering thresholds.

| Nanoparticles | Nanoparticles_tubulin | Nanoparticles_vim2 | Nanoparticles_actin |
| --- | --- | --- | --- |
| 384 | 306 | 405 | 181 |

To visualize gene expression patterns, a heatmap was constructed using the 100 most differentially expressed genes (Figure 2). Genes were sorted by variability, and a Z-score of normalized expression was calculated for each sample.

**Figure 2.**
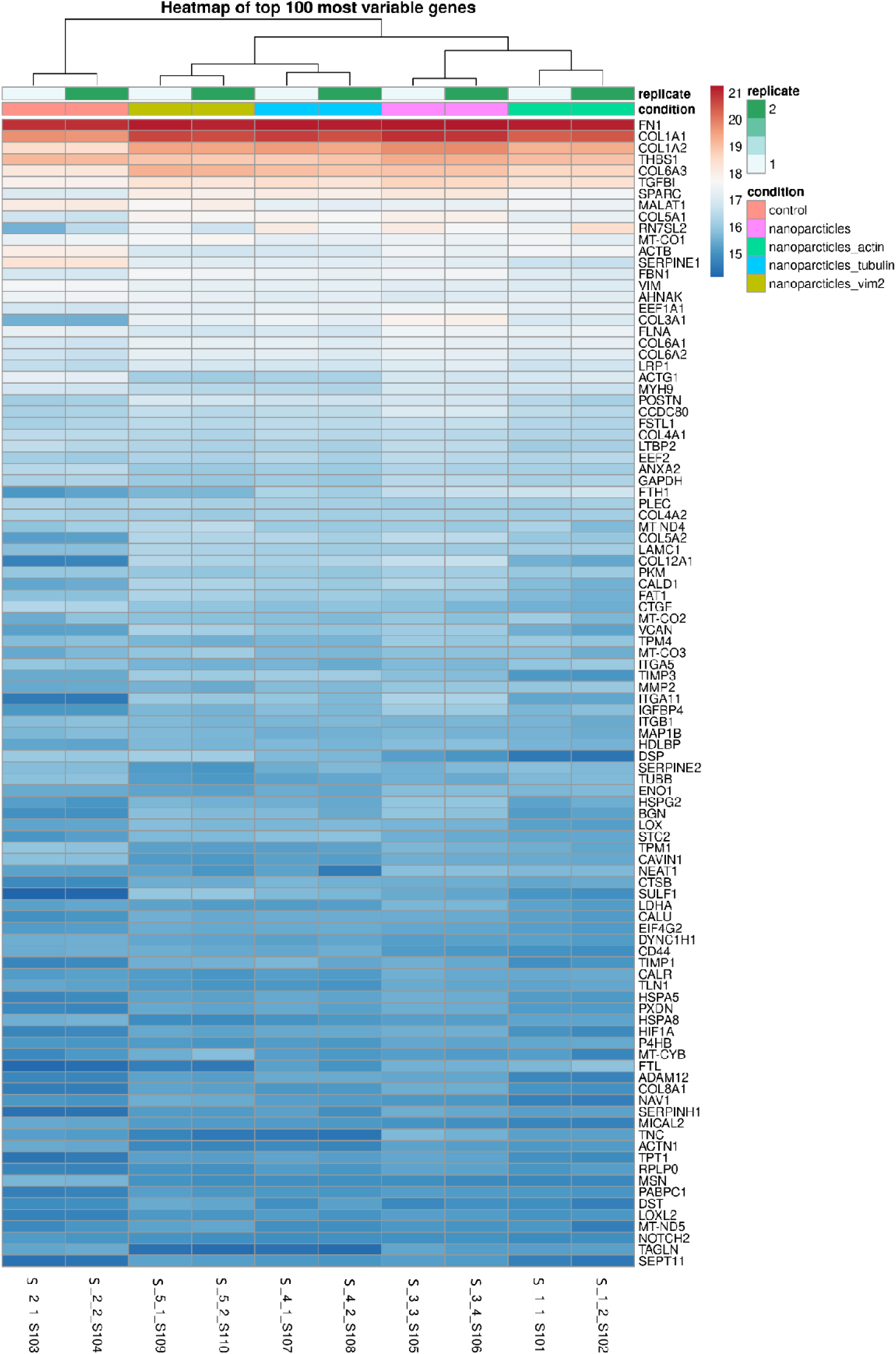
Top 100 of the most variable genes.

The samples clustered by experimental groups, indicating a correlation between experimental conditions on the transcriptomic profiles of cells. All experimental groups form separate clusters, with the nanoparticles_tubulin and nanoparticles_vim2 groups clustering closer together than with nanoparticles_actin, which may reflect biological similarities between tubulin- and vimentin-related changes.

Among the most variable genes, extracellular matrix and adhesion genes predominated: FN1, COL1A1, COL1A2, COL6A1, COL6A2, LAMC1, SPARC, FBN1. Their high variability suggests a profound remodeling of cell–matrix interactions.

Genes associated with cytoskeleton reorganization were also identified, such as VIM (vimentin), TUBB (beta-tubulin), ACTN1 (alpha-actinin-1), and MYH9 (myosin heavy chain 9), which suggests the presence of feedback mechanisms within interference with cytoskeleton. Additionally, stress-associated genes (HSP90, HSPB1, HSPG2) showed activity, indicating a stress response triggered by the presence of nanoparticles inside the cells.

Volcano plots were designed for all experimental groups to assess the effect of nanoparticles on gene expression (nanoparticles, nanoparticles_tubulin, nanoparticles_vim2, and nanoparticles_actin) (Figure 3).

**Figure 3.**
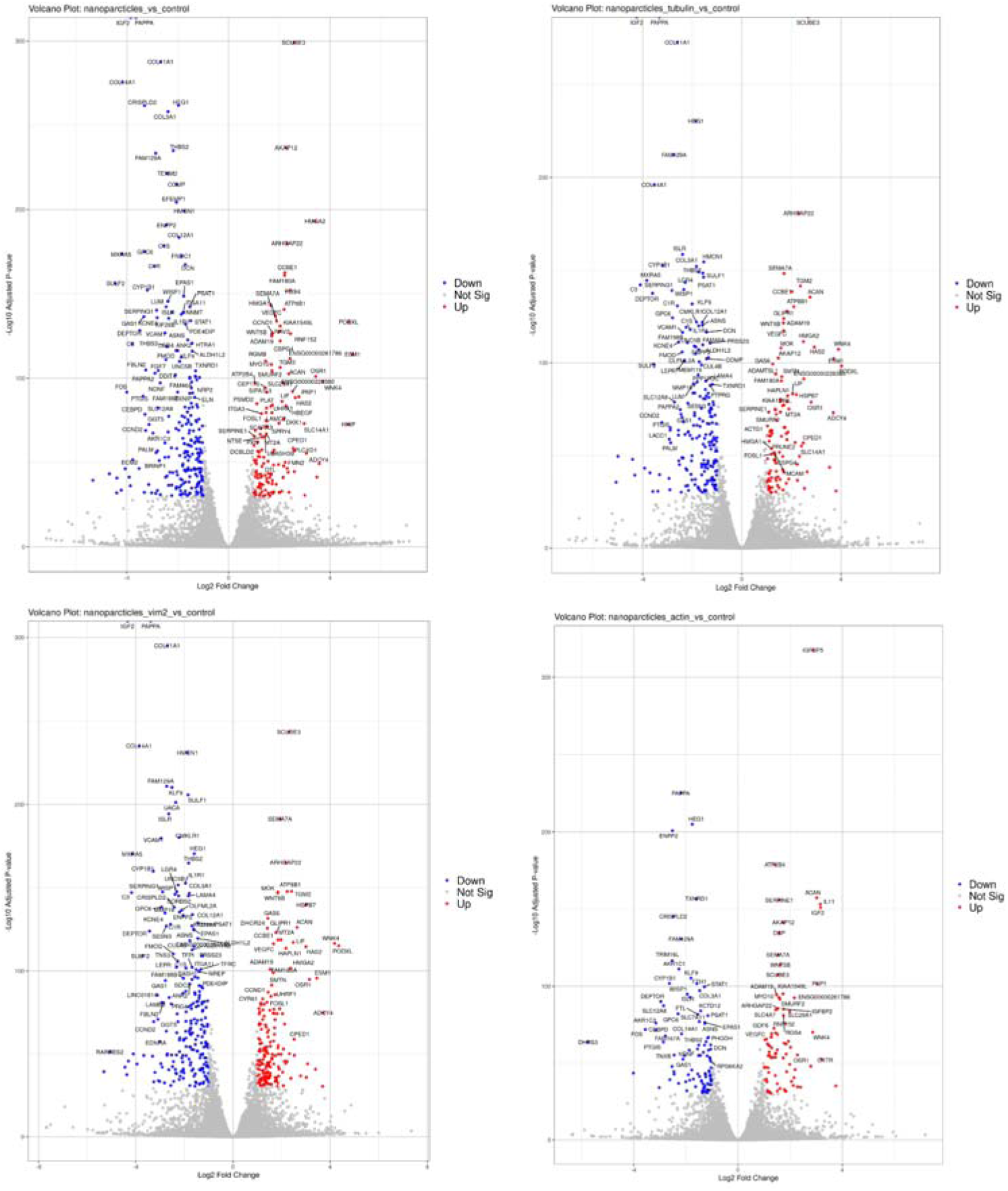
Volcano-plots of DEG distribution between groups.

The pattern of changes in up- and down-regulated genes varied significantly depending on the type of cytoskeletal modification. In the nanoparticles group, activation of oxidative stress-related genes (e.g., SOD2) and suppression of mitochondrial components (COX4I1, CYC1) were detected, suggesting the mitochondrial dysfunction under nanoparticle exposure. In the nanoparticles_tubulin group, activation of mitotic cycle genes (ANLN, KIF11) was observed, possibly indicating disruption of cell division processes due to nanoparticle effects on the mitotic spindle. The nanoparticles_vim2 group exhibited predominant changes in genes associated with extracellular matrix remodeling (SERPINE1, MMP2), reflecting potential alterations in cell–environment interactions. In the nanoparticles_actin group, activation of actin structure reorganization genes (ACTG1, PFN1) and suppression of cell adhesion molecules (CDH1, EPCAM) were detected, suggesting a potential loss of intercellular contacts. Overall, there was a consistent link between cytoskeletal interference and changes in cytoskeleton-associated gene expression.

A comparative intersection analysis (UpSet plot) revealed 98 common DEGs shared across all experimental groups, along with unique DEG sets for each: 82 for nanoparticles, 7 for nanoparticles_tubulin, 98 for nanoparticles_vim2, 33 for nanoparticles_actin, and 3 for the nanoparticle–protein crosslinked groups (Figure 4 and Table 2).

**Figure 4.**
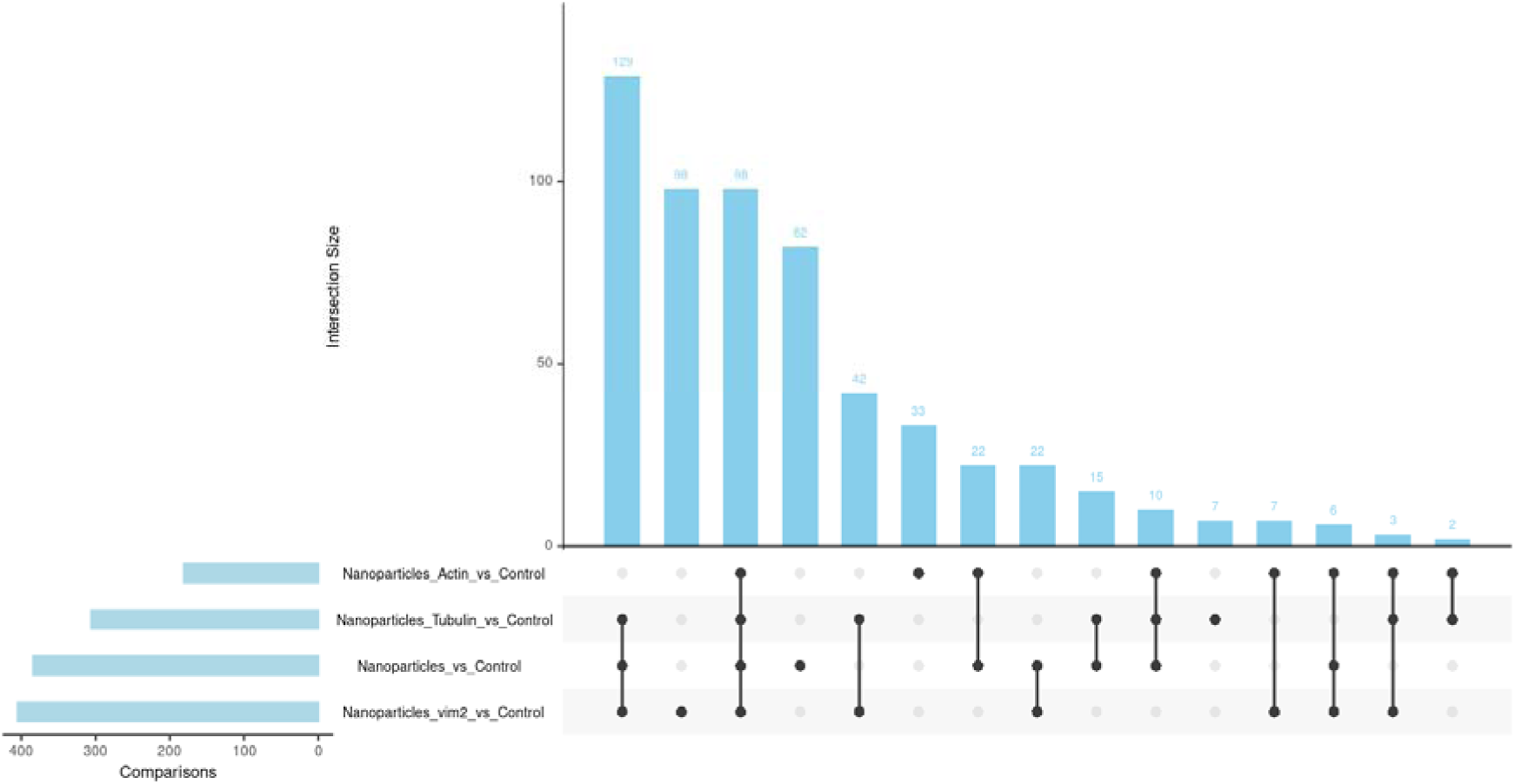
UpSet-plot of common and unique DEGs in the study groups.

**Table 2.**
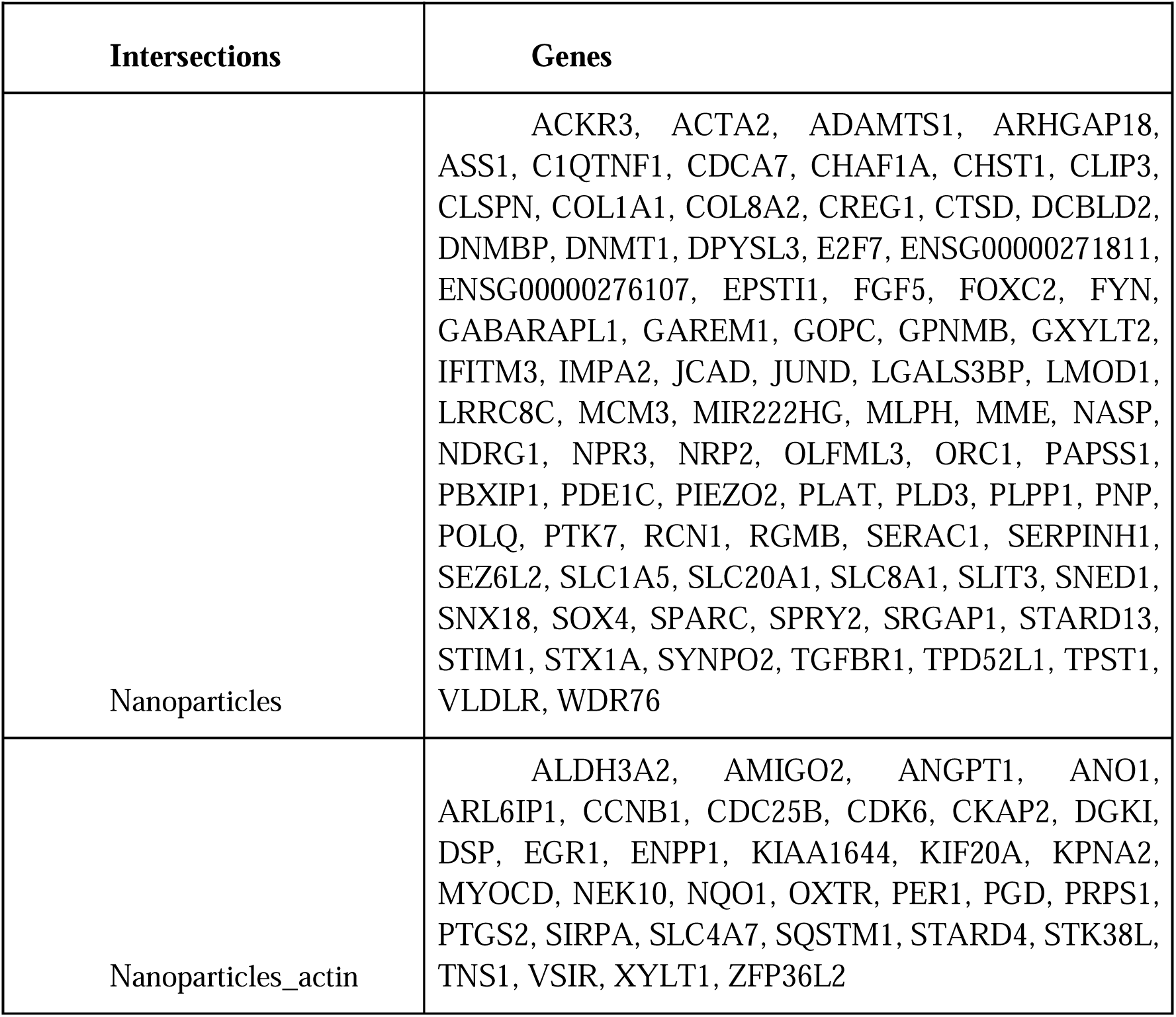
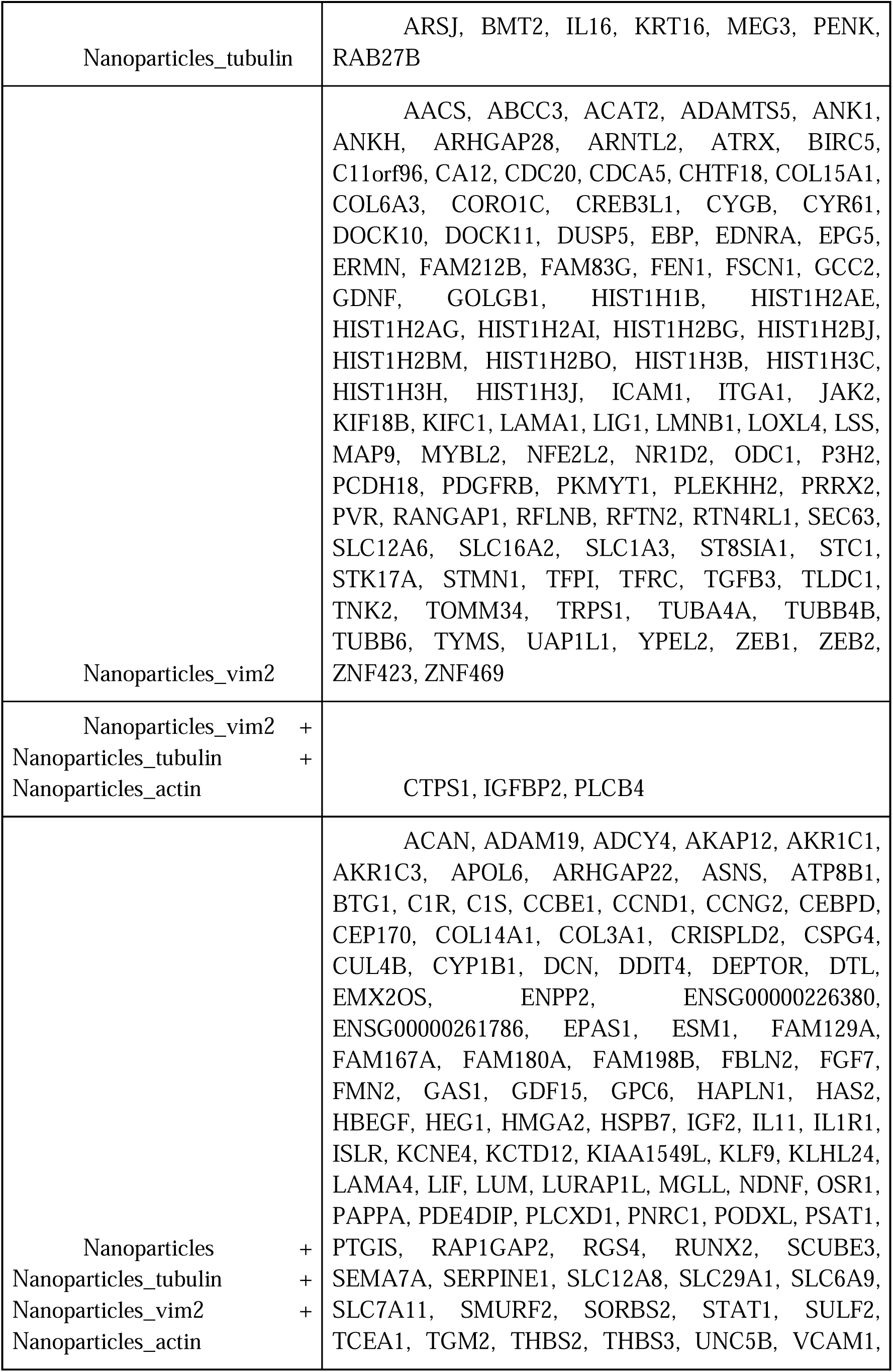
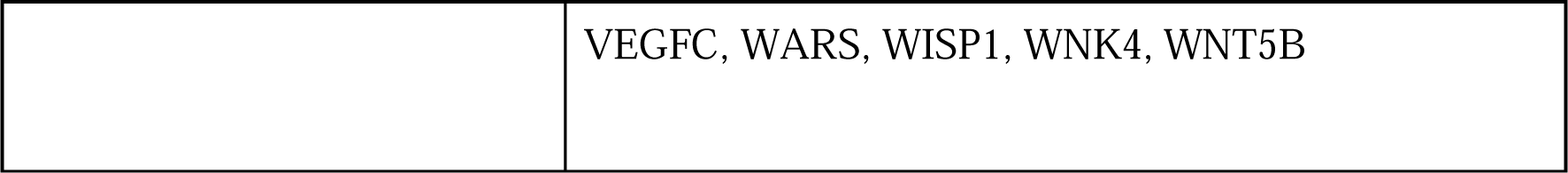
Commonly expressed genes identified across and within the experimental groups.

### 3.3 Investigation of Cytoskeleton-Associated Genes

To explore how cytoskeleton-related genes respond to nanoparticle exposure, we constructed a heatmap of the 100 most variable cytoskeleton-associated genes (Figure 5). A complete list of cytoskeleton-associated genes can be found in <u>Supplementary 2</u>. Interestingly, the analysis revealed two major clusters that separated the samples into *control* and *nanoparticles_actin*, and *nanoparticles*, *nanoparticles_vim2*, and *nanoparticles_tubulin*. It was also noted that the composition of the top 100 general DEGs and the top 100 cytoskeleton-specific DEGs differed significantly, likely highlighting the specificity of cytoskeletal alterations.

**Figure 5.**
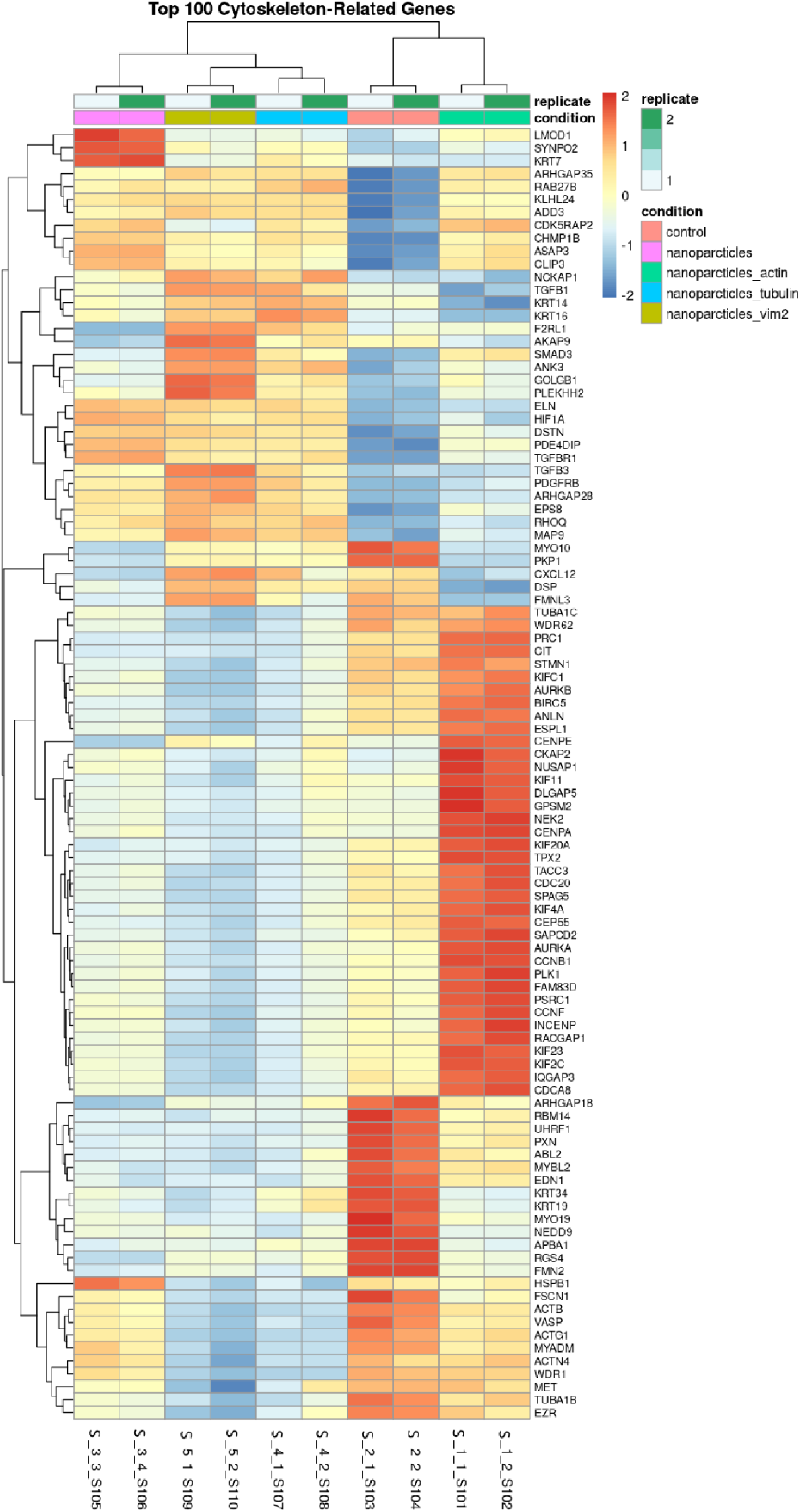
Heatmap of the 100 most variable cytoskeleton-related genes.

A notable upregulation was observed for genes involved in spindle formation and cell division, including *KIF11*, *CENPA*, *AURKB*, *KIF23*, and *CDC20*. This suggests the activation of proliferative pathways and potential induction of cytoskeletal reorganization in response to nanoparticle interference with structural proteins.

Interestingly, samples treated with actin-targeted nanoparticles displayed a distinct expression profile that partially overlapped with both control samples and other cytoskeleton-targeted groups. This suggests that nanoparticles with affinity for actin may provoke a more pronounced transcriptional response.

Overall, these findings indicate that targeted action to nanoparticles with affinity for specific cytoskeletal elements triggers distinct and robust transcriptional programs, particularly enhancing pathways associated with mitotic spindle dynamics, cell cycle progression, and cytoskeleton remodeling.

An UpSet plot revealed the presence of 4 shared DEGs across all studied groups and unique sets of DEGs for each: 7 for *nanoparticles*, 2 for *nanoparticles_tubulin*, 12 for *nanoparticles_vim2*, and 4 for *nanoparticles_actin* (Figure 6 and Table 3).

**Figure 6.**
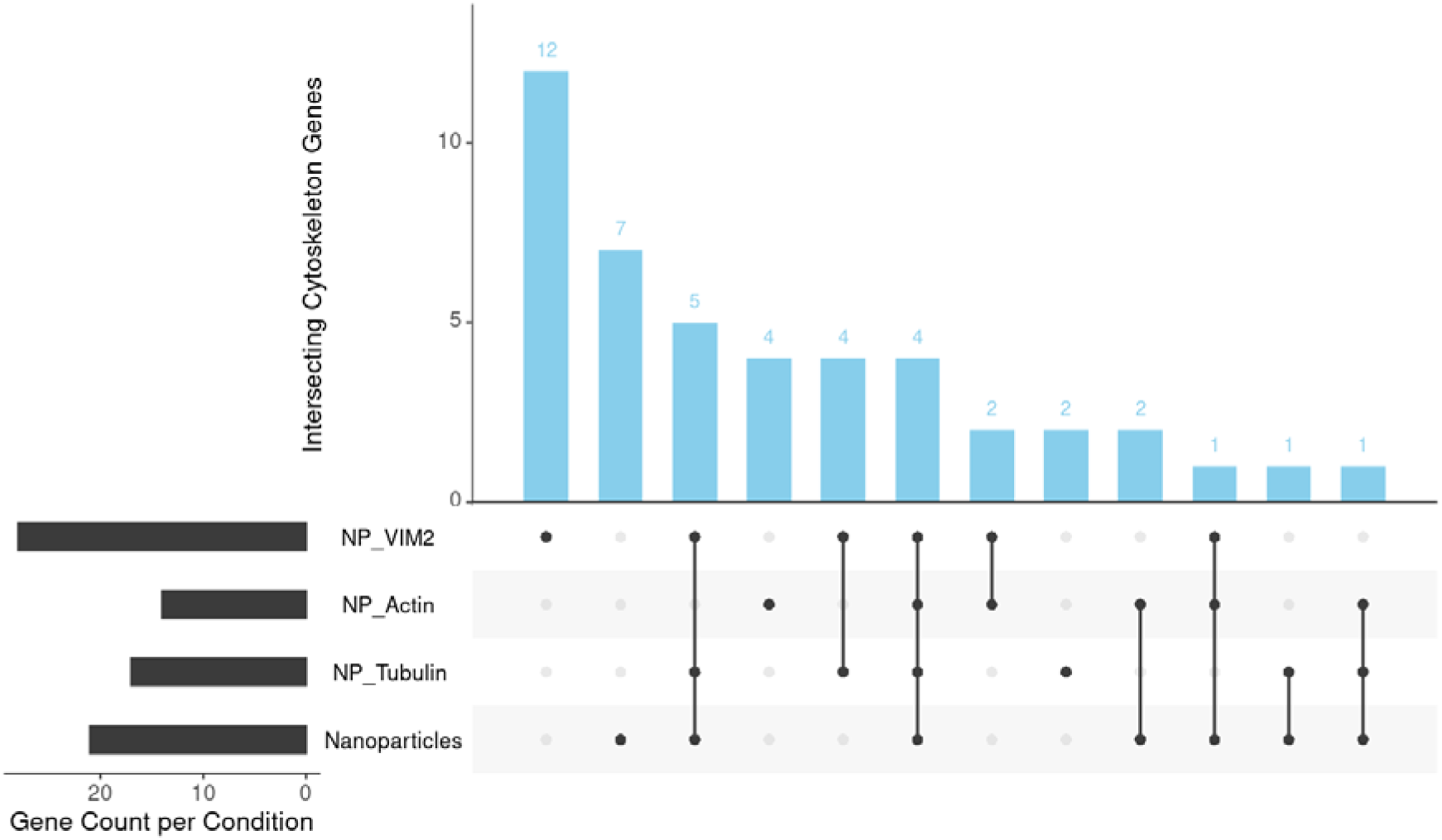
UpSet-plot of common and unique cytoskeleton DEGs in the study groups.

**Table 3.**
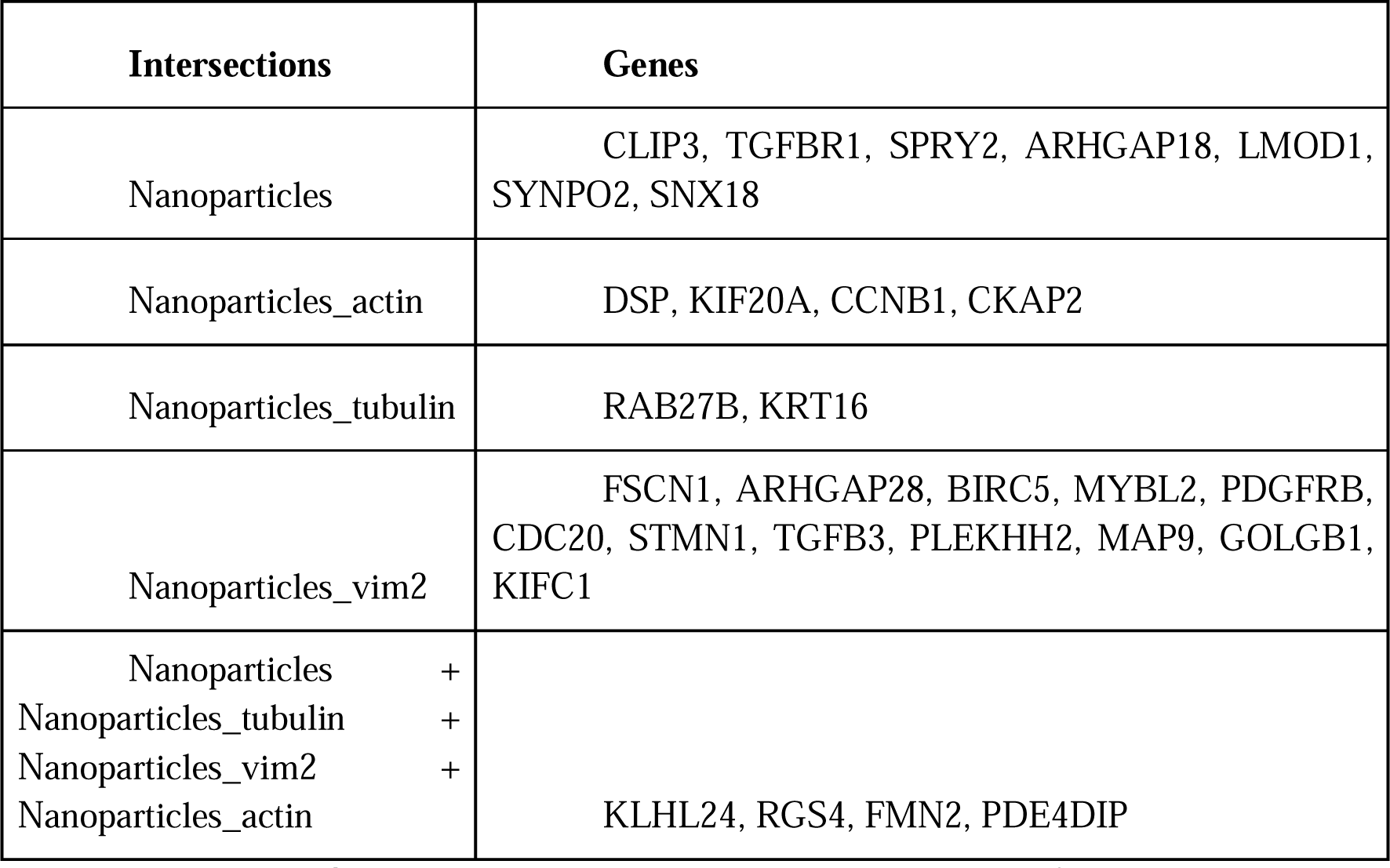
Cytoskeleton-associated genes identified in each group and intersections between groups.

The *nanoparticles* group included specific genes such as *CLIP3*, *TGFBR1*, *SPRY2*, *ARHGAP18*, *LMOD1*, *SYNPO2*, and *SNX18*, which are involved in regulating cell adhesion, growth signaling pathways, cytoskeletal reorganization, and early stress responses to nanoparticle internalization. These genes may participate in early or general responses to nanoparticle exposure.

In the *nanoparticles_actin* group, four genes — *DSP*, *KIF20A*, *CCNB1*, and *CKAP2* — were associated with cell cycle control, mitotic coordination, and maintenance of cytoskeletal integrity, reflecting the activation of proliferative and structural responses to actin-mediated mechanical stress. The *nanoparticles_tubulin* group included two key genes, *RAB27B* and *KRT16*, involved in regulating cellular architecture and microtubule dynamics. The *nanoparticles_vim2* group had a broader set of 12 genes (*FSCN1*, *ARHGAP28*, *BIRC5*, *MYBL2*, *PDGFRB*, *CDC20*, *STMN1*, *TGFB3*, *PLEKHH2*, *MAP9*, *GOLGB1*, and *KIFC1*) that regulate processes related to cell proliferation, actin cytoskeleton remodeling, mitotic events, and reorganization of intercellular interactions, indicating activation of proliferative and migratory programs in response to intermediate filament deformation.

The intersection of all groups included four genes (*KLHL24*, *RGS4*, *FMN2*, and *PDE4DIP*), which may reflect a universal transcriptional response to cytoskeletal damage or tension, participating in the regulation of actin dynamics, signaling, and intracellular architecture organization.

Thus, the data demonstrate both specific and overlapping molecular changes in response to different types of nanoparticles, impacting the structure and function of the cytoskeleton. These findings formed the basis for further functional annotations through GO, KEGG, and GSEA analyses.

### 3.4 Gene Ontology (GO) Analysis of Biological Processes

Genes with differential expression statistics exceeding the ad hoc threshold (-log10 (p value) > 30, |log2 (Fold Change)| > 1) were subjected to Gene Ontology (GO) analysis using the clusterProfiler R package (v. 4.8.2). The “Biological Processes” database was used for enrichment analysis. The results are presented in Figure 7.

**Figure 7.**
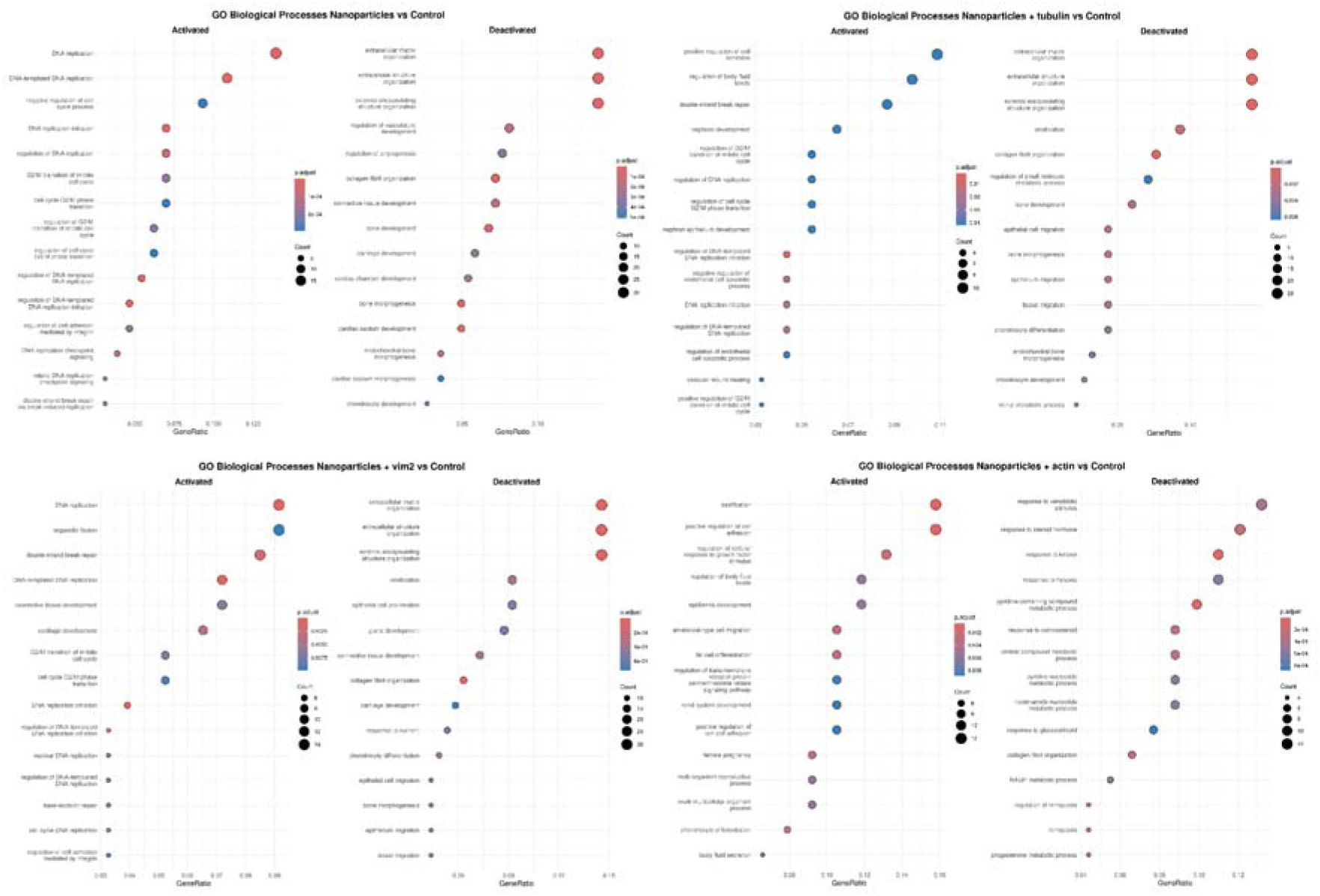
Results of Gene Ontology analysis of groups.

Functional annotation of differentially expressed genes (DEG) using Gene Ontology (GO) revealed enrichment of the following biological processes common across all investigated groups (Table 4):

**Table 4.** - Commonly Activated and Deactivated Biological Processes Across All Investigated Groups.

| Commonly Activated Biological Processes | Commonly Deactivated Biological Processes |
| --- | --- |
| G2/M transition of mitotic cell cycle<br>regulation of G2/M transition of mitotic cell cycle<br>regulation of cell cycle G2/M phase transition<br>nephron development | extracellular matrix organization<br>extracellular structure organization<br>external encapsulating structure organization<br>collagen fibril organization<br>bone morphogenesis |
| nephron epithelium development<br>regulation of body fluid levels | bone development<br>connective tissue development<br>endochondral bone morphogenesis<br>cartilage development<br>chondrocyte differentiation<br>respiratory system development<br>epithelial cell proliferation<br>organ growth<br>regulation of epithelial cell proliferation<br>cellular response to corticosteroid stimulus<br>transforming growth factor beta receptor superfamily signaling pathway<br>glial cell-derived neurotrophic factor receptor signaling pathway<br>response to nutrient<br>retinal metabolic process<br>cell-substrate adhesion<br>aldehyde metabolic process |

Table 4 presents the list of biological processes that were commonly activated or deactivated in all experimental groups. The most prominent processes are related to mitotic division and the cell cycle, including: *G2/M transition of mitotic cell cycle*, *regulation of G2/M transition of mitotic cell cycle*, and *regulation of cell cycle G2/M phase transition*. This indicates regulation of cell division or enhanced control at the critical G2/M checkpoint, where the cell decides whether to proceed into mitosis. Such a response is often associated with repair, stress, or proliferation as a reaction to cellular damage.

The extensive list of suppressed processes relates to organ and tissue development (*nephron development*, *bone morphogenesis*, *cartilage development*, *connective tissue development*, *chondrocyte differentiation*, *respiratory system development*). This suggests inhibition of differentiation and morphogenesis, particularly in tissues of mesenchymal and epithelial origin. Likely, the cells are “abandoning” complex developmental programs in favor of an emergency stress response.

Structural and extracellular processes are also suppressed (*extracellular matrix organization*, *collagen fibril organization*, *cell-substrate adhesion*), which may indicate disruptions in intercellular interactions, tissue architecture, and potentially cell migration. This is typical following exposure to toxic agents, including nanoparticles, when cells lose adhesion and extracellular organization.

Some hormonal and signaling pathways are suppressed as well (*transforming growth factor beta receptor superfamily signaling pathway*, *cellular response to corticosteroid stimulus*), reflecting inhibition of anti-stress and anti-inflammatory activities, as well as impaired transmission of growth and immune signals. Suppression of pathways involved in metabolism and nutrition (*response to nutrient*, *retinal metabolic process*, *aldehyde metabolic process*) may point to a metabolic shift, decreased energy metabolism or adaptation to limited resources/toxins.

Overall, nanoparticles induce a universal activation of the cell cycle, likely as a compensatory mechanism following damage, while simultaneously suppressing key developmental, morphogenetic, intercellular, and hormonal regulatory processes. This indicates cellular stress, disruption of homeostasis, and a possible shift toward pathogenesis if exposure persists or intensifies.

To gain deeper insights into the molecular mechanisms underlying the observed biological processes, KEGG pathway analysis was subsequently conducted to identify the signaling pathways involved.

### 3.5 KEGG Pathway Analysis

To identify disrupted molecular mechanisms under the influence of nanoparticles, KEGG (Kyoto Encyclopedia of Genes and Genomes) pathway enrichment analysis was performed for each of the studied groups: *nanoparticles*, *nanoparticles_tubulin*, *nanoparticles_vim2*, and *nanoparticles_actin*. The analysis allowed the identification of both activated and suppressed pathways (Figure 8), as well as the visualization of gene interactions through network graphs.

**Figure 8.**
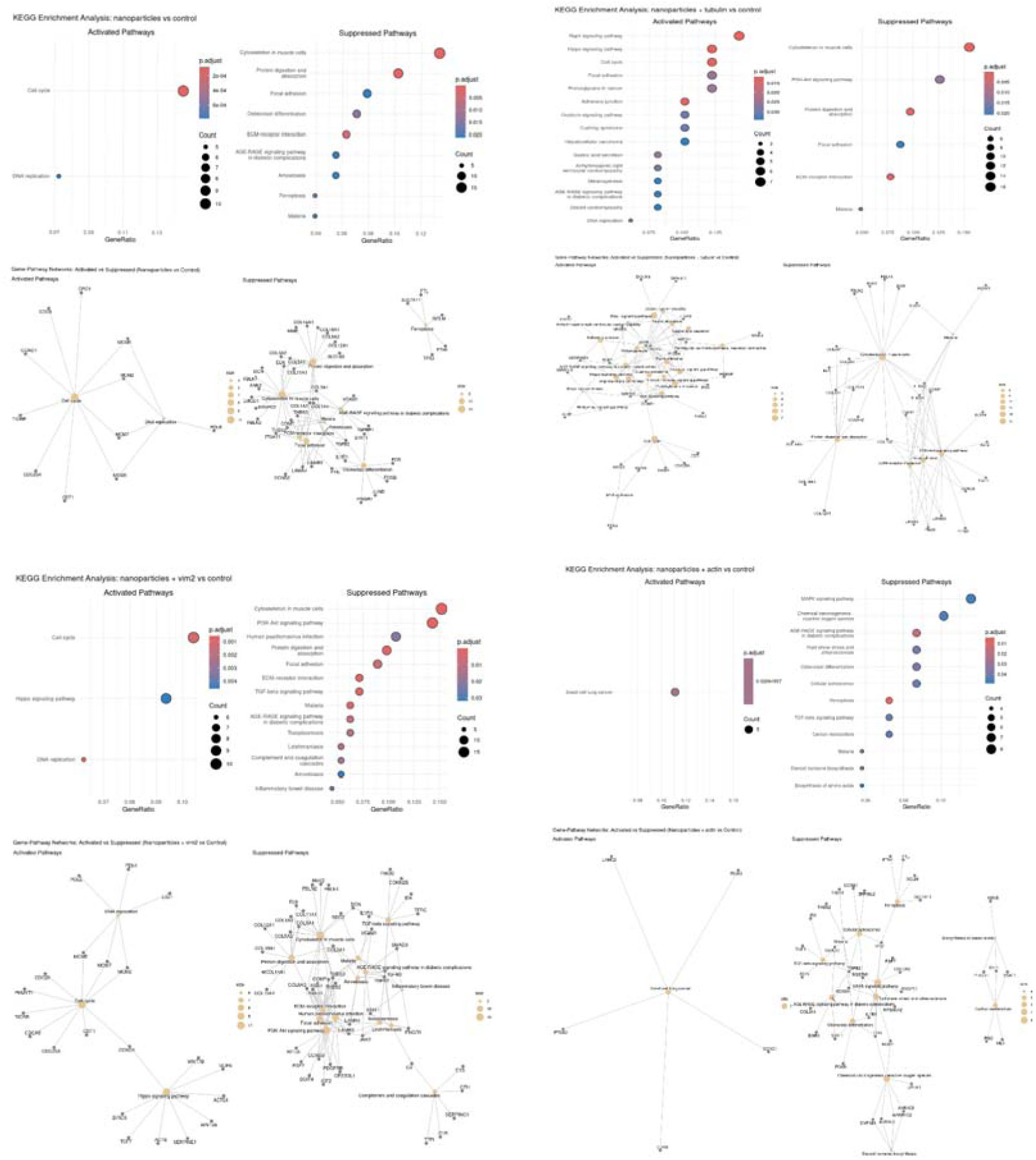
Results of KEGG analysis of signaling pathways and genes of the experimental groups.

A list of common activated and deactivated pathways is presented in Table 5. Table 5 provides a summary of signaling pathways that were either activated or deactivated across all experimental conditions (nanoparticles, nanoparticles_tubulin, nanoparticles_vim2, nanoparticles_actin) based on the KEGG analysis results. Among the activated pathways, cascades related to the regulation of cell growth and division predominated, such as the Hippo signaling pathway, Cushing syndrome, Hedgehog signaling pathway, and Proteoglycans in cancer, as well as pathways controlling morphogenesis (Melanogenesis) and cellular structure (Cytoskeleton in muscle cells).

**Table 5.** - Common activated and deactivated signaling pathways for all studied groups according to KEGG (Kyoto Encyclopedia of Genes and Genomes)

| Common activated signaling pathways | Common deactivated signaling pathways |
| --- | --- |
| Hippo signaling pathway<br>Wnt signaling pathway<br>Cushing syndrome<br>Melanogenesis<br>Hedgehog signaling pathway<br>Focal adhesion<br>Proteoglycans in cancer | Cytoskeleton in muscle cells<br>ECM-receptor interaction<br>Amoebiasis<br>Malaria<br>PI3K-Akt signaling pathway<br>p53 signaling pathway<br>Pertussis |

Among the common deactivated pathways, key signaling systems responsible for cell adhesion, intercellular interactions, and immune responses were identified: Wnt signaling pathway, Focal adhesion, ECM-receptor interaction, PI3K-Akt signaling pathway, and p53 signaling pathway. The suppression of these pathways suggests disruption of extracellular matrix integrity, reduced cellular survival, and impaired stress response regulation mechanisms.

Additionally, among the deactivated cascades, pathways associated with infectious and inflammatory processes (Amoebiasis, Malaria, Pertussis) were noted, which aligns with the observed decrease in immune gene activity under cytoskeletal damage.

In the *nanoparticles* group, significant enrichment of cell cycle pathways was identified, including the activation of genes such as *CDK1*, *CCNB1*, *CCNA2*, and *CDC20*, indicating the initiation of mitotic processes. This is further supported by the inclusion of the DNA replication pathway, involving *MCM* complex genes (*MCM2–MCM7*) and *PCNA*. Conversely, suppression of the PI3K-Akt, TGF-beta, ECM-receptor interaction, and Focal adhesion signaling cascades was observed, where key components include *COL1A1*, *FN1*, *ITGA5*, and *TGFBR2*. These changes may indicate weakening of cell adhesion, structural integrity, and reparative signaling transmission. The network graph showed that *COL1A1*, *THBS1*, and *FN1* act as key nodes, participating in multiple pathways and forming critical points of nanoparticle impact on cellular functions.

In the *nanoparticles_tubulin* group, a similar pattern of cell cycle and DNA replication pathway activation was observed, involving *CDK1*, *BUB1*, and the *MCM* complex. Suppression of the Focal adhesion, PI3K-Akt, and MAPK pathways was accompanied by decreased expression of *COL4A2*, *ITGB1*, *BRAF*, and *RAF1*. Interestingly, suppression of cascades related to intercellular signaling, such as the neuroactive ligand-receptor interaction and calcium signaling pathway, was also observed, suggesting disturbances in calcium homeostasis and intracellular signaling. Genes *THBS1*, *ANGPT2*, and *ITGA5* formed dense hubs of suppressed interactions in the graphs, highlighting the vulnerability of cellular communications under microtubule architecture disruption.

In the *nanoparticles_vim2* group, activation of cell cycle genes and the Hippo signaling pathway was observed, which can be interpreted as a compensatory cellular response to stress and disruption of intermediate filament integrity. Vimentin plays a crucial role in maintaining mechanical stability, and its dysfunction can reduce the strength of the cellular framework. In response, cells likely activate division and growth programs (through *YAP1*, *TEAD1*, and suppression of *LATS2*) to enhance regenerative processes, maintain cell numbers, and adapt to mechanical changes. Activation of the Hippo signaling pathway reflects an attempt to control cell size and proliferation: when the cytoskeleton is damaged, cells may activate *YAP1* to stimulate survival and growth. Suppression of *LATS2*, which inhibits *YAP1*, further enhances this proliferative stimulus.

On the other hand, significant suppression of innate immunity and inflammatory signaling genes (such as *CCL2*, IL6, *CXCL10*, *TLR2*, *ICAM1*) indicates weakened defensive functions. This may be due to disrupted mechanical and inflammatory signal transmission to the nucleus following intermediate filament damage, as vimentin participates in the presentation of innate immunity receptors on the cell surface. Its disruption leads to reduced recognition of danger-associated molecular patterns (PAMPs) and immune activation.

Furthermore, suppression of numerous immune genes functioning across multiple pathways (e.g., *CXCL8*, *TNFSF10*, *IL1R1*) suggests a systemic weakening of innate immunity, making cells less capable of detecting stressors and coordinating inflammatory responses. This could be a protective strategy to avoid excessive inflammation when the cytoskeleton is compromised or a manifestation of impaired signaling due to the loss of structural integrity.

In the *nanoparticles_actin* group, the smallest number of activated pathways was observed. Enrichment was detected only for the small cell lung cancer pathway, involving proliferative genes *MYC* and *BCL2*. At the same time, strong suppression of pathways regulating cell survival, proliferation, and interactions with the environment, including PI3K-Akt, MAPK, and Rap1 signaling cascades (involving *AKT1*, *EGFR*, *ITGA2*, and *MAPK1*), was noted. Additionally, significant suppression of metabolic pathways (glycolysis, fatty acid, and amino acid metabolism) was observed due to decreased activity of *LDHA*, *PFKP*, and *ACADL*. Network analysis confirmed the suppression of both signaling and metabolic hubs, with *VEGFA*, *IGF1*, and *MAPK3* occupying central positions in the graphs and participating in multiple suppressed pathways, forming modules of complex impact.

Collectively, the data show that the most universal cellular response to nanoparticle exposure is the activation of division and replication programs, whereas suppression predominantly affects signaling cascades, immune responses, and structural components of the cell surface. Specific effects are further enhanced or modified depending on the cytoskeletal element present under the exposure.

The most pronounced suppression of inflammatory and metabolic cascades was observed in the presence of actin, whereas in the case of vimentin, disruptions were mainly observed in immune receptor systems and inflammatory signaling transmission. The KEGG analysis results complement the GO analysis and highlight the complex, multi-layered cellular response to nanoparticles, combining elements of proliferation activation, structural destabilization, and suppression of intercellular communications.

### 3.6 Pathway Enrichment Analysis (FGSEA)

Pathway analysis using the FGSEA method for hallmark gene sets revealed significant enrichment of cellular proliferative programs under nanoparticle exposure and their combinations with cytoskeletal modulation (Figure 9, Supplementary 3).

**Figure 9.**
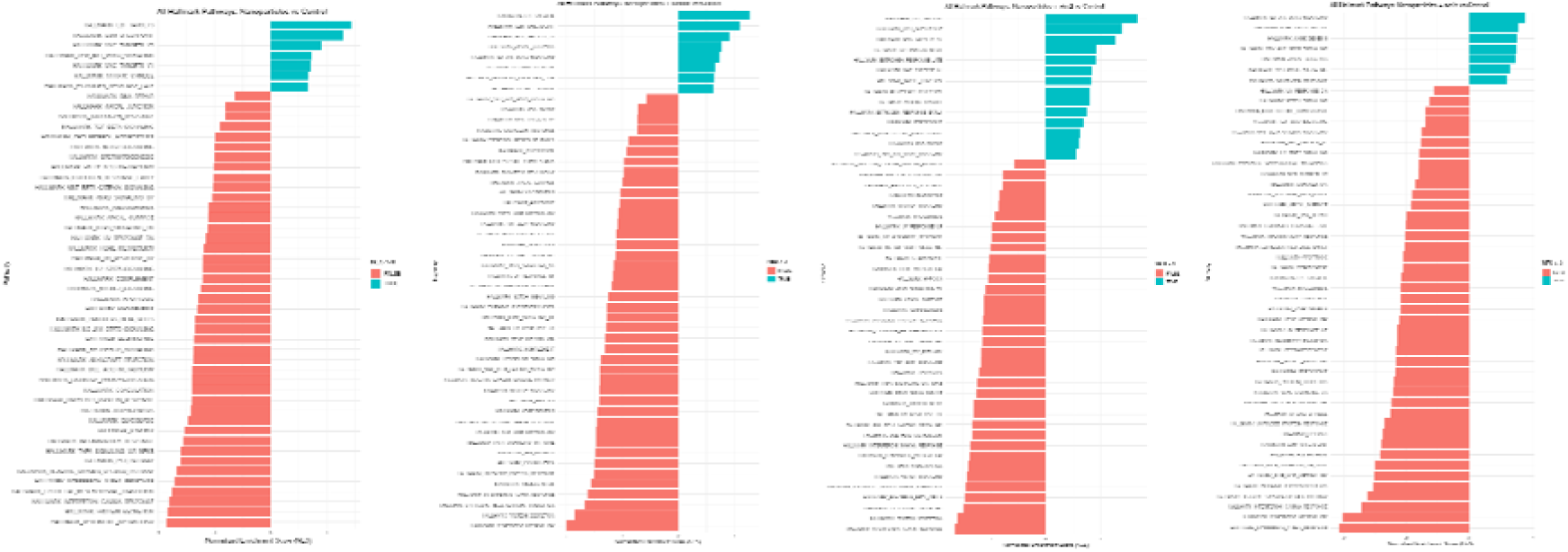
Results of FGSEA analysis of the study groups.

In the *nanoparticles* condition, the E2F Targets (NES = 1.43, padj = 0.0209) and G2M Checkpoint (NES = 1.29, padj = 0.0545) pathways were significantly enriched, indicating enhancement of cell cycle processes. Among the leading-edge genes involved in these pathways were *HMGA1*, *CDC25A*, *PSMC3IP*, *E2F2*, and *CCND1*, which play key roles in transcription regulation and mitotic progression. This may reflect a proliferative response of the cells to nanoparticle exposure.

Under the *nanoparticles_tubulin* condition, the activation trend of E2F Targets (NES = 1.27, padj = 0.072) and G2M Checkpoint (NES = 1.10, padj = 0.39) pathways persisted, although the statistical significance was reduced. The leading-edge genes included *CDC25A*, *CENPM*, *PSMC3IP*, *MT2A*, and *E2F2*, also suggesting partial activity of the mitotic response. However, the weaker statistical support may indicate a change in the signaling background due to disruption of microtubule structure.

The most pronounced and statistically significant enrichment was observed in the *nanoparticles_vim 2* group, where the E2F Targets pathway demonstrated the highest NES = 1.67 with padj = 0.0053. The G2M Checkpoint pathway was also significantly enriched (NES = 1.38, padj = 0.042). Leading genes such as *CDC25A*, *CENPM*, *AURKB*, *E2F2*, *MT2A*, and *PLK4* were more strongly activated compared to other groups. Such an enhancement of proliferative programs may be related to the destabilization of intermediate filaments, leading to cellular remodeling and the reinitiation of cell division.

The *nanoparticles_actin* group showed no significant enrichment of hallmark pathways. All presented signaling cascades, including IL6/JAK/STAT3 signaling, KRAS signaling up, and Angiogenesis, had high padj values (1.0) and low NES (< 0.9), indicating a lack of statistical significance. Nevertheless, leading-edge genes such as *IL9R*, *IL2RG*, *CNTFR*, *ABCB1*, *OLR1*, *PRG2*, and *LPL*, involved in the regulation of immune and angiogenic processes, were identified, which may reflect compensatory or stress responses to mechanical disruption of the actin network.

Overall, the FGSEA results demonstrate that proliferative signaling pathways are most sensitive to nanoparticle exposure combined with vimentin network disruption, whereas interference with the actin cytoskeleton does not induce significant enrichment of hallmark programs. This highlights the key role of intermediate filaments in regulating cellular proliferation and sensitivity to external signals.

The transcriptomic analysis showed that exposure to free nanoparticles, as well as conjugates with cytoskeletal elements, induces pronounced and reproducible changes in gene expression, encompassing both general cellular stress responses and specific programs related to cytoskeletal dynamics, cell cycle, and extracellular matrix interactions. These results suggest that targeting actin, tubulin, and vimentin structures triggers distinct transcriptional scenarios, reflecting the functional specialization of the corresponding cytoskeletal components. These findings provide a foundation for further functional analyses and discussion of the mechanisms underlying the observed effects.

## 4. Discussion

In this study, we investigated the effect of magnetically controlled mechanical tension on gene expression patterns in human mesenchymal stem cells (MSCs, bone marrow-derived fibroblasts). Using RNA sequencing followed by differential expression analysis, Gene Ontology (GO) enrichment, Kyoto Encyclopedia of Genes and Genomes (KEGG) pathway analysis, and principal component analysis (PCA), we characterized the transcriptional response of MSCs to mechanical stimulation mediated by magnetic nanoparticle systems.

Our results demonstrate a profound remodeling of the transcriptional landscape under the influence of mechanical tension on cytoskeletal elements. Differential expression analysis revealed a substantial number of up- and down-regulated genes, with volcano plots and PCA clearly showing a clear separation between mechanically stimulated and control groups. Notably, genes related to cytoskeletal organization, cell cycle regulation, and extracellular matrix (ECM) remodeling were markedly affected.

GO enrichment analysis indicated that mechanical tension predominantly influences biological processes related to cytoskeletal organization (e.g., “actin filament organization,” “microtubule-based process”) and cell division (e.g., “mitotic spindle organization,” “chromosome segregation”). These findings align with previous observations that mechanical stimuli can modulate cytoskeletal architecture and cell proliferation in stem cells.

KEGG pathway analysis further highlighted the activation of pathways associated with the cell cycle, DNA replication, and mitotic processes. For example, the significant regulation of key regulators such as *CDC20*, *CCNB1*, *CDK1*, and *AURKB* points to enhanced proliferative activity in response to mechanical stimulation. This observation is consistent with the concept of stem cell fate regulation driven by mechanotransduction, where cytoskeletal tension promotes cell cycle progression and expansion [10.1016/j.tcb.2022.03.010].

Conversely, we observed a marked suppression of several signaling and metabolic pathways. Downregulated pathways included those involved in modulating immune responses (“TNF signaling pathway,” “NF-κB signaling pathway”) as well as metabolic cascades such as oxidative phosphorylation and glycolysis/gluconeogenesis. These results suggest a shift in cellular priorities under mechanical tension, favoring proliferation and cytoskeletal reorganization over immune reactivity and metabolic activity. Suppression of inflammatory signaling pathways may also contribute to a more “immune silent” microenvironment, potentially enhancing the regenerative potential of MSCs under mechanical conditioning. Additionally, the presence of iron oxide nanoparticles within mesenchymal stem cells could independently trigger the activation of anti-inflammatory pathways [16].

Interestingly, cytoskeleton-associated pathways such as “regulation of actin cytoskeleton” and “focal adhesion” exhibited a complex pattern, with activation of certain regulatory genes (e.g., *RHOA*, *ROCK1*, *ACTN1*) alongside suppression of other modulators (e.g., *PTK2/FAK*, *VCL*). This dual modulation likely reflects dynamic remodeling of focal adhesion complexes and cytoskeletal tension necessary for mechanosensing and adaptation to mechanical stimuli. Suppression of *FAK* may indicate destabilization of actin filaments; given that FAK activation inhibits Rho signaling at sites of actin filament assembly [17], it is possible that interference with actin cytoskeletal elements instead promotes activation of these pathways.

Network analysis provided further insights into the relationships between differentially expressed genes. Highly interconnected nodes involved in mitotic control (e.g., *BUB1*, *PLK1*, *TOP2A*) and cytoskeletal dynamics (e.g., *MYH9*, *FLNA*, *TUBB*) formed prominent hubs, suggesting that mechanical tension induces a coordinated transcriptional program to support cytoskeletal stability and cell cycle progression.

Collectively, our data indicate that magnetically induced mechanical tension elicits a strong proliferative response in MSCs while simultaneously modulating cytoskeletal organization and suppressing immune and metabolic pathways. These findings provide valuable insights into how mechanical forces shape stem cell behavior and open new opportunities for optimizing mechanical conditioning strategies to enhance MSC-based regenerative therapies.

## 5. Conclusion

Understanding the role of individual cytoskeletal elements brings us closer to a better comprehension of human cell physiology and advances our knowledge of the causes behind pathological processes. The observed physiological responses of cells to the presence of cytoskeleton-associated nanoparticles are partly fundamental in nature, representing a reaction to foreign inclusions within the cells. At the same time, it is clearly evident that cells exhibit distinct biological responses to the deformation of specific cytoskeletal elements induced by magnetic nanoparticles in a magnetic field. The obtained results open new opportunities for non-invasive investigation of intracellular processes in living cells over time.

Future studies should further explore the long-term effects of mechanical tension on the differentiation potential and aging of mesenchymal stem cells (MSCs), as well as analyze the mechanotransduction pathways responsible for mediating the observed changes in gene expression. Additionally, integrating proteomic and metabolomic approaches will be crucial for validating the transcriptional findings at the functional level.

## 6. Acknowledgments

The reported study was funded by the Russian Science Foundation Grant #22-74-10041.

We are grateful to IOS UB RAS (Theme No. 124020500044-4) in part of Fe3O4 nanoparticles synthesis and analysis, and the work and characterisation of Fe@C nanoparticles was performed within the framework of the state assignment of the Ministry of Education and Science of Russia for IFM UB RAS

## Notes

https://drive.google.com/drive/folders/1pKDPhBjhLcgYJVeLvb5PR-9VS_ZMxAni?usp=drive_link

https://drive.google.com/file/d/1-HJ-wnmXF_Ia4q2Hhg5Scu5IDIGKvrP3/view?usp=drive_link

https://drive.google.com/file/d/1s4u0IwifmnWrppW0xkoAxZ92-vT3I3Mp/view?usp=drive_link

### Competing Interest Statement

The authors have declared no competing interest.

