## Supplementary figures and images for "Magnetically controlled tension of cytoskeletal elements with magnetic nanoparticles affects the expression of signaling pathway genes associated with cytoskeletal elements"

### Supplementary 2

#### Top 100 Cytoskeleton-Related Genes

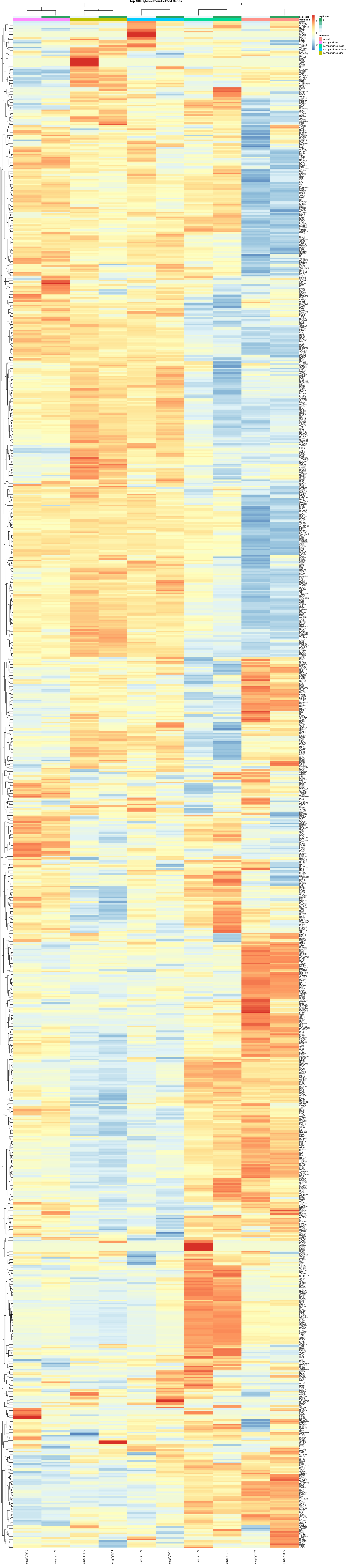
