## Supplementary 3 for "Magnetically controlled tension of cytoskeletal elements with magnetic nanoparticles affects the expression of signaling pathway genes associated with cytoskeletal elements"

### Nanoparticles

| Pathway | NES | pval | padj |
|---|---|---|---|
| HALLMARK_E2F_TARGETS | 1.43 | $7.1\cdot10^{-4}$ | $2.1\cdot10^{-2}$ |
| HALLMARK_G2M_CHECKPOINT | 1.29 | $2.5\cdot10^{-2}$ | $5.5\cdot10^{-2}$ |
| HALLMARK_MYC_TARGETS_V2 | 0.90 | $6.7\cdot10^{-1}$ | $7.9\cdot10^{-1}$ |
| HALLMARK_PI3K_AKT_MTOR_SIGNALING | 0.72 | $9.4\cdot10^{-1}$ | $1.0\cdot10^{0}$ |
| HALLMARK_MYC_TARGETS_V1 | 0.71 | $9.8\cdot10^{-1}$ | $1.0\cdot10^{0}$ |
| HALLMARK_MITOTIC_SPINDLE | 0.67 | $9.9\cdot10^{-1}$ | $1.0\cdot10^{0}$ |
| HALLMARK_ESTROGEN_RESPONSE_LATE | 0.66 | $9.9\cdot10^{-1}$ | $1.0\cdot10^{0}$ |
| HALLMARK_DNA_REPAIR | -0.65 | $1.0\cdot10^{0}$ | $1.0\cdot10^{0}$ |
| HALLMARK_APICAL_JUNCTION | -0.80 | $1.0\cdot10^{0}$ | $1.0\cdot10^{0}$ |
| HALLMARK_ANDROGEN_RESPONSE | -0.81 | $9.7\cdot10^{-1}$ | $1.0\cdot10^{0}$ |
| HALLMARK_TGF_BETA_SIGNALING | -0.91 | $7.0\cdot10^{-1}$ | $8.1\cdot10^{-1}$ |
| HALLMARK_CHOLESTEROL_HOMEOSTASIS | -0.99 | $4.9\cdot10^{-1}$ | $5.9\cdot10^{-1}$ |
| HALLMARK_NOTCH_SIGNALING | -1.00 | $4.3\cdot10^{-1}$ | $5.5\cdot10^{-1}$ |
| HALLMARK_SPERMATOGENESIS | -1.00 | $4.4\cdot10^{-1}$ | $5.5\cdot10^{-1}$ |
| HALLMARK_FATTY_ACID_METABOLISM | -1.01 | $4.2\cdot10^{-1}$ | $5.5\cdot10^{-1}$ |
| HALLMARK_ESTROGEN_RESPONSE_EARLY | -1.02 | $3.2\cdot10^{-1}$ | $4.5\cdot10^{-1}$ |
| HALLMARK_WNT_BETA_CATENIN_SIGNALING | -1.02 | $3.8\cdot10^{-1}$ | $5.2\cdot10^{-1}$ |
| HALLMARK_KRAS_SIGNALING_UP | -1.04 | $3.3\cdot10^{-1}$ | $4.6\cdot10^{-1}$ |
| HALLMARK_ANGIOGENESIS | -1.11 | $2.6\cdot10^{-1}$ | $3.8\cdot10^{-1}$ |
| HALLMARK_APICAL_SURFACE | -1.11 | $2.5\cdot10^{-1}$ | $3.7\cdot10^{-1}$ |
| HALLMARK_KRAS_SIGNALING_DN | -1.13 | $1.0\cdot10^{-1}$ | $1.6\cdot10^{-1}$ |
| HALLMARK_UV_RESPONSE_DN | -1.15 | $7.6\cdot10^{-2}$ | $1.3\cdot10^{-1}$ |
| HALLMARK_HEME_METABOLISM | -1.19 | $3.9\cdot10^{-2}$ | $7.0\cdot10^{-2}$ |
| HALLMARK_UV_RESPONSE_UP | -1.21 | $2.2\cdot10^{-2}$ | $5.1\cdot10^{-2}$ |
| HALLMARK_IL2_STAT5_SIGNALING | -1.21 | $3.3\cdot10^{-2}$ | $6.1\cdot10^{-2}$ |
| HALLMARK_COMPLEMENT | -1.21 | $3.2\cdot10^{-2}$ | $6.1\cdot10^{-2}$ |
| HALLMARK_MTORC1_SIGNALING | -1.24 | $2.6\cdot10^{-2}$ | $5.5\cdot10^{-2}$ |
| HALLMARK_APOPTOSIS | -1.29 | $1.4\cdot10^{-2}$ | $3.5\cdot10^{-2}$ |
| HALLMARK_MYOGENESIS | -1.31 | $2.1\cdot10^{-2}$ | $5.1\cdot10^{-2}$ |
| HALLMARK_PANCREAS_BETA_CELLS | -1.34 | $8.2\cdot10^{-2}$ | $1.3\cdot10^{-1}$ |
| HALLMARK_IL6_JAK_STAT3_SIGNALING | -1.34 | $2.8\cdot10^{-2}$ | $5.6\cdot10^{-2}$ |
| HALLMARK_PEROXISOME | -1.36 | $9.0\cdot10^{-3}$ | $2.8\cdot10^{-2}$ |
| HALLMARK_HEDGEHOG_SIGNALING | -1.37 | $5.5\cdot10^{-2}$ | $9.5\cdot10^{-2}$ |
| HALLMARK_ALLOGRAFT_REJECTION | -1.38 | $1.2\cdot10^{-2}$ | $3.3\cdot10^{-2}$ |
| HALLMARK_BILE_ACID_METABOLISM | -1.39 | $8.0\cdot10^{-3}$ | $2.7\cdot10^{-2}$ |
| HALLMARK_OXIDATIVE_PHOSPHORYLATION | -1.40 | $9.8\cdot10^{-3}$ | $2.9\cdot10^{-2}$ |
| HALLMARK_COAGULATION | -1.40 | $6.2\cdot10^{-3}$ | $2.2\cdot10^{-2}$ |
| HALLMARK_UNFOLDED_PROTEIN_RESPONSE | -1.40 | $1.1\cdot10^{-2}$ | $2.9\cdot10^{-2}$ |
| HALLMARK_ADIPOGENESIS | -1.41 | $5.3\cdot10^{-3}$ | $2.1\cdot10^{-2}$ |
| HALLMARK_GLYCOLYSIS | -1.47 | $5.3\cdot10^{-3}$ | $2.1\cdot10^{-2}$ |
| HALLMARK_HYPOXIA | -1.53 | $5.3\cdot10^{-3}$ | $2.1\cdot10^{-2}$ |
| HALLMARK_INFLAMMATORY_RESPONSE | -1.55 | $5.1\cdot10^{-3}$ | $2.1\cdot10^{-2}$ |
| HALLMARK_TNFA_SIGNALING_VIA_NFKB | -1.59 | $5.3\cdot10^{-3}$ | $2.1\cdot10^{-2}$ |
| HALLMARK_P53_PATHWAY | -1.60 | $5.3\cdot10^{-3}$ | $2.1\cdot10^{-2}$ |
| HALLMARK_REACTIVE_OXYGEN_SPECIES_PATHWAY | -1.68 | $2.2\cdot10^{-3}$ | $2.1\cdot10^{-2}$ |
| HALLMARK_INTERFERON_ALPHA_RESPONSE | -1.71 | $3.0\cdot10^{-3}$ | $2.1\cdot10^{-2}$ |
| HALLMARK_EPITHELIAL_MESENCHYMAL_TRANSITION | -1.76 | $5.4\cdot10^{-3}$ | $2.1\cdot10^{-2}$ |
| HALLMARK_INTERFERON_GAMMA_RESPONSE | -1.80 | $5.1\cdot10^{-3}$ | $2.1\cdot10^{-2}$ |
| HALLMARK_PROTEIN_SECRETION | -1.84 | $1.5\cdot10^{-3}$ | $2.1\cdot10^{-2}$ |
| HALLMARK_XENOBIOTIC_METABOLISM | -1.85 | $4.6\cdot10^{-3}$ | $2.1\cdot10^{-2}$ |

### Nanoparticles + tubulin

| Pathway | NES | pval | padj |
|---|---|---|---|
| HALLMARK_E2F_TARGETS | 1.27 | $3.0\cdot10^{-2}$ | $7.2\cdot10^{-2}$ |
| HALLMARK_G2M_CHECKPOINT | 1.10 | $2.6\cdot10^{-1}$ | $3.9\cdot10^{-1}$ |
| HALLMARK_MYC_TARGETS_V2 | 0.90 | $6.5\cdot10^{-1}$ | $8.4\cdot10^{-1}$ |
| HALLMARK_APICAL_JUNCTION | 0.77 | $9.4\cdot10^{-1}$ | $1.0\cdot10^{0}$ |
| HALLMARK_IL6_JAK_STAT3_SIGNALING | 0.73 | $9.3\cdot10^{-1}$ | $1.0\cdot10^{0}$ |
| HALLMARK_SPERMATOGENESIS | 0.65 | $9.9\cdot10^{-1}$ | $1.0\cdot10^{0}$ |
| HALLMARK_ESTROGEN_RESPONSE_LATE | 0.64 | $10.0\cdot10^{-1}$ | $1.0\cdot10^{0}$ |
| HALLMARK_MITOTIC_SPINDLE | 0.63 | $10.0\cdot10^{-1}$ | $1.0\cdot10^{0}$ |
| HALLMARK_PI3K_AKT_MTOR_SIGNALING | -0.57 | $1.0\cdot10^{0}$ | $1.0\cdot10^{0}$ |
| HALLMARK_DNA_REPAIR | -0.72 | $1.0\cdot10^{0}$ | $1.0\cdot10^{0}$ |
| HALLMARK_MYC_TARGETS_V1 | -0.72 | $1.0\cdot10^{0}$ | $1.0\cdot10^{0}$ |
| HALLMARK_ANDROGEN_RESPONSE | -0.75 | $9.9\cdot10^{-1}$ | $1.0\cdot10^{0}$ |
| HALLMARK_ESTROGEN_RESPONSE_EARLY | -0.89 | $9.4\cdot10^{-1}$ | $1.0\cdot10^{0}$ |
| HALLMARK_PEROXISOME | -0.92 | $7.3\cdot10^{-1}$ | $9.1\cdot10^{-1}$ |
| HALLMARK_PANCREAS_BETA_CELLS | -0.96 | $5.1\cdot10^{-1}$ | $6.9\cdot10^{-1}$ |
| HALLMARK_CHOLESTEROL_HOMEOSTASIS | -0.96 | $5.7\cdot10^{-1}$ | $7.4\cdot10^{-1}$ |
| HALLMARK_APICAL_SURFACE | -1.00 | $4.5\cdot10^{-1}$ | $6.2\cdot10^{-1}$ |
| HALLMARK_MYOGENESIS | -1.02 | $4.0\cdot10^{-1}$ | $5.7\cdot10^{-1}$ |
| HALLMARK_APOPTOSIS | -1.05 | $2.7\cdot10^{-1}$ | $3.9\cdot10^{-1}$ |
| HALLMARK_FATTY_ACID_METABOLISM | -1.07 | $2.1\cdot10^{-1}$ | $3.4\cdot10^{-1}$ |
| HALLMARK_TGF_BETA_SIGNALING | -1.09 | $2.6\cdot10^{-1}$ | $3.9\cdot10^{-1}$ |
| HALLMARK_GLYCOLYSIS | -1.10 | $1.2\cdot10^{-1}$ | $2.3\cdot10^{-1}$ |
| HALLMARK_ALLOGRAFT_REJECTION | -1.10 | $1.6\cdot10^{-1}$ | $2.7\cdot10^{-1}$ |
| HALLMARK_IL2_STAT5_SIGNALING | -1.10 | $1.4\cdot10^{-1}$ | $2.4\cdot10^{-1}$ |
| HALLMARK_KRAS_SIGNALING_DN | -1.15 | $7.0\cdot10^{-2}$ | $1.3\cdot10^{-1}$ |
| HALLMARK_UV_RESPONSE_UP | -1.16 | $5.0\cdot10^{-2}$ | $1.0\cdot10^{-1}$ |
| HALLMARK_INFLAMMATORY_RESPONSE | -1.18 | $3.4\cdot10^{-2}$ | $7.8\cdot10^{-2}$ |
| HALLMARK_NOTCH_SIGNALING | -1.23 | $1.4\cdot10^{-1}$ | $2.4\cdot10^{-1}$ |
| HALLMARK_OXIDATIVE_PHOSPHORYLATION | -1.25 | $1.7\cdot10^{-2}$ | $4.7\cdot10^{-2}$ |
| HALLMARK_KRAS_SIGNALING_UP | -1.27 | $2.2\cdot10^{-2}$ | $5.7\cdot10^{-2}$ |
| HALLMARK_HEME_METABOLISM | -1.29 | $1.1\cdot10^{-2}$ | $3.7\cdot10^{-2}$ |
| HALLMARK_UV_RESPONSE_DN | -1.29 | $9.1\cdot10^{-3}$ | $3.3\cdot10^{-2}$ |
| HALLMARK_COMPLEMENT | -1.30 | $1.2\cdot10^{-2}$ | $3.7\cdot10^{-2}$ |
| HALLMARK_HEDGEHOG_SIGNALING | -1.37 | $5.7\cdot10^{-2}$ | $1.1\cdot10^{-1}$ |
| HALLMARK_WNT_BETA_CATENIN_SIGNALING | -1.40 | $4.0\cdot10^{-2}$ | $8.8\cdot10^{-2}$ |
| LLMARK_REACTIVE_OXYGEN_SPECIES_PATHWAY | -1.40 | $2.9\cdot10^{-2}$ | $7.2\cdot10^{-2}$ |
| HALLMARK_MTORC1_SIGNALING | -1.40 | $6.4\cdot10^{-3}$ | $2.6\cdot10^{-2}$ |
| HALLMARK_HYPOXIA | -1.40 | $6.4\cdot10^{-3}$ | $2.6\cdot10^{-2}$ |
| HALLMARK_ADIPOGENESIS | -1.43 | $6.7\cdot10^{-3}$ | $2.6\cdot10^{-2}$ |
| HALLMARK_INTERFERON_GAMMA_RESPONSE | -1.43 | $5.7\cdot10^{-3}$ | $2.6\cdot10^{-2}$ |
| HALLMARK_TNFA_SIGNALING_VIA_NFKB | -1.45 | $6.7\cdot10^{-3}$ | $2.6\cdot10^{-2}$ |
| HALLMARK_P53_PATHWAY | -1.47 | $6.4\cdot10^{-3}$ | $2.6\cdot10^{-2}$ |
| HALLMARK_BILE_ACID_METABOLISM | -1.47 | $3.4\cdot10^{-3}$ | $2.6\cdot10^{-2}$ |
| HALLMARK_COAGULATION | -1.52 | $2.3\cdot10^{-3}$ | $2.6\cdot10^{-2}$ |
| HALLMARK_UNFOLDED_PROTEIN_RESPONSE | -1.52 | $1.9\cdot10^{-3}$ | $2.6\cdot10^{-2}$ |
| HALLMARK_ANGIOGENESIS | -1.53 | $1.6\cdot10^{-2}$ | $4.7\cdot10^{-2}$ |
| HALLMARK_INTERFERON_ALPHA_RESPONSE | -1.63 | $1.6\cdot10^{-3}$ | $2.6\cdot10^{-2}$ |
| MARK_EPITHELIAL_MESENCHYMAL_TRANSITION | -1.64 | $6.8\cdot10^{-3}$ | $2.6\cdot10^{-2}$ |
| HALLMARK_PROTEIN_SECRETION | -1.86 | $1.6\cdot10^{-3}$ | $2.6\cdot10^{-2}$ |
| HALLMARK_XENOBIOTIC_METABOLISM | -1.96 | $5.6\cdot10^{-3}$ | $2.6\cdot10^{-2}$ |

### Nanoparticles + vim2

| Pathway | NES | pval | padj |
|---|---|---|---|
| HALLMARK_E2F_TARGETS | 1.67 | $1.1\cdot10^{-4}$ | $5.3\cdot10^{-3}$ |
| HALLMARK_G2M_CHECKPOINT | 1.38 | $5.7\cdot10^{-3}$ | $4.2\cdot10^{-2}$ |
| HALLMARK_MYC_TARGETS_V2 | 1.27 | $1.1\cdot10^{-1}$ | $2.4\cdot10^{-1}$ |
| HALLMARK_SPERMATOGENESIS | 0.92 | $6.4\cdot10^{-1}$ | $9.4\cdot10^{-1}$ |
| HALLMARK_ESTROGEN_RESPONSE_LATE | 0.92 | $6.9\cdot10^{-1}$ | $9.8\cdot10^{-1}$ |
| HALLMARK_MYC_TARGETS_V1 | 0.85 | $8.4\cdot10^{-1}$ | $10.0\cdot10^{-1}$ |
| HALLMARK_APICAL_JUNCTION | 0.83 | $8.6\cdot10^{-1}$ | $10.0\cdot10^{-1}$ |
| HALLMARK_ALLOGRAFT_REJECTION | 0.80 | $9.0\cdot10^{-1}$ | $10.0\cdot10^{-1}$ |
| HALLMARK_MITOTIC_SPINDLE | 0.80 | $9.1\cdot10^{-1}$ | $10.0\cdot10^{-1}$ |
| HALLMARK_ESTROGEN_RESPONSE_EARLY | 0.75 | $9.6\cdot10^{-1}$ | $10.0\cdot10^{-1}$ |
| HALLMARK_PEROXISOME | 0.69 | $9.7\cdot10^{-1}$ | $10.0\cdot10^{-1}$ |
| HALLMARK_CHOLESTEROL_HOMEOSTASIS | 0.63 | $9.9\cdot10^{-1}$ | $10.0\cdot10^{-1}$ |
| HALLMARK_DNA_REPAIR | 0.60 | $10.0\cdot10^{-1}$ | $10.0\cdot10^{-1}$ |
| HALLMARK_PI3K_AKT_MTOR_SIGNALING | 0.56 | $10.0\cdot10^{-1}$ | $10.0\cdot10^{-1}$ |
| LLMARK_REACTIVE_OXYGEN_SPECIES_PATHWAY | -0.58 | $10.0\cdot10^{-1}$ | $10.0\cdot10^{-1}$ |
| HALLMARK_FATTY_ACID_METABOLISM | -0.78 | $9.9\cdot10^{-1}$ | $10.0\cdot10^{-1}$ |
| HALLMARK_ANDROGEN_RESPONSE | -0.78 | $9.7\cdot10^{-1}$ | $10.0\cdot10^{-1}$ |
| HALLMARK_GLYCOLYSIS | -0.84 | $9.8\cdot10^{-1}$ | $10.0\cdot10^{-1}$ |
| HALLMARK_MTORC1_SIGNALING | -0.86 | $9.5\cdot10^{-1}$ | $10.0\cdot10^{-1}$ |
| HALLMARK_MYOGENESIS | -0.92 | $7.6\cdot10^{-1}$ | $10.0\cdot10^{-1}$ |
| HALLMARK_UV_RESPONSE_UP | -0.99 | $4.8\cdot10^{-1}$ | $7.6\cdot10^{-1}$ |
| HALLMARK_INFLAMMATORY_RESPONSE | -0.99 | $5.0\cdot10^{-1}$ | $7.6\cdot10^{-1}$ |
| HALLMARK_IL6_JAK_STAT3_SIGNALING | -1.01 | $4.2\cdot10^{-1}$ | $6.7\cdot10^{-1}$ |
| HALLMARK_COMPLEMENT | -1.02 | $3.5\cdot10^{-1}$ | $5.9\cdot10^{-1}$ |
| HALLMARK_HEME_METABOLISM | -1.04 | $2.9\cdot10^{-1}$ | $5.0\cdot10^{-1}$ |
| HALLMARK_HYPOXIA | -1.05 | $2.7\cdot10^{-1}$ | $4.8\cdot10^{-1}$ |
| HALLMARK_KRAS_SIGNALING_DN | -1.10 | $1.6\cdot10^{-1}$ | $3.0\cdot10^{-1}$ |
| HALLMARK_APICAL_SURFACE | -1.12 | $2.4\cdot10^{-1}$ | $4.5\cdot10^{-1}$ |
| HALLMARK_ADIPOGENESIS | -1.13 | $1.1\cdot10^{-1}$ | $2.4\cdot10^{-1}$ |
| HALLMARK_UNFOLDED_PROTEIN_RESPONSE | -1.15 | $1.4\cdot10^{-1}$ | $2.9\cdot10^{-1}$ |
| HALLMARK_OXIDATIVE_PHOSPHORYLATION | -1.15 | $8.6\cdot10^{-2}$ | $2.2\cdot10^{-1}$ |
| HALLMARK_IL2_STAT5_SIGNALING | -1.16 | $8.2\cdot10^{-2}$ | $2.2\cdot10^{-1}$ |
| HALLMARK_P53_PATHWAY | -1.18 | $5.8\cdot10^{-2}$ | $1.7\cdot10^{-1}$ |
| HALLMARK_TGF_BETA_SIGNALING | -1.18 | $1.5\cdot10^{-1}$ | $3.0\cdot10^{-1}$ |
| HALLMARK_APOPTOSIS | -1.21 | $4.7\cdot10^{-2}$ | $1.5\cdot10^{-1}$ |
| HALLMARK_TNFA_SIGNALING_VIA_NFKB | -1.24 | $2.6\cdot10^{-2}$ | $9.9\cdot10^{-2}$ |
| HALLMARK_KRAS_SIGNALING_UP | -1.27 | $2.3\cdot10^{-2}$ | $9.9\cdot10^{-2}$ |
| HALLMARK_ANGIOGENESIS | -1.31 | $9.0\cdot10^{-2}$ | $2.2\cdot10^{-1}$ |
| HALLMARK_UV_RESPONSE_DN | -1.31 | $2.0\cdot10^{-2}$ | $9.9\cdot10^{-2}$ |
| HALLMARK_WNT_BETA_CATENIN_SIGNALING | -1.34 | $6.6\cdot10^{-2}$ | $1.8\cdot10^{-1}$ |
| HALLMARK_BILE_ACID_METABOLISM | -1.34 | $2.4\cdot10^{-2}$ | $9.9\cdot10^{-2}$ |
| HALLMARK_INTERFERON_GAMMA_RESPONSE | -1.37 | $6.4\cdot10^{-3}$ | $4.2\cdot10^{-2}$ |
| HALLMARK_XENOBIOTIC_METABOLISM | -1.39 | $4.7\cdot10^{-3}$ | $4.2\cdot10^{-2}$ |
| HALLMARK_COAGULATION | -1.41 | $6.7\cdot10^{-3}$ | $4.2\cdot10^{-2}$ |
| HALLMARK_NOTCH_SIGNALING | -1.44 | $3.9\cdot10^{-2}$ | $1.3\cdot10^{-1}$ |
| MARK_EPITHELIAL_MESENCHYMAL_TRANSITION | -1.46 | $1.8\cdot10^{-3}$ | $2.2\cdot10^{-2}$ |
| HALLMARK_PANCREAS_BETA_CELLS | -1.47 | $3.9\cdot10^{-2}$ | $1.3\cdot10^{-1}$ |
| HALLMARK_HEDGEHOG_SIGNALING | -1.50 | $2.0\cdot10^{-2}$ | $9.9\cdot10^{-2}$ |
| HALLMARK_PROTEIN_SECRETION | -1.61 | $7.5\cdot10^{-4}$ | $1.2\cdot10^{-2}$ |
| HALLMARK_INTERFERON_ALPHA_RESPONSE | -1.67 | $7.5\cdot10^{-4}$ | $1.2\cdot10^{-2}$ |

### Nanoparticles + actin

| Pathway | NES | pval | padj |
|---|---|---|---|
| HALLMARK_IL6_JAK_STAT3_SIGNALING | 0.86 | $7.4\cdot10^{-1}$ | $1.0\cdot10^{0}$ |
| HALLMARK_KRAS_SIGNALING_UP | 0.75 | $9.5\cdot10^{-1}$ | $1.0\cdot10^{0}$ |
| HALLMARK_ANGIOGENESIS | 0.74 | $8.7\cdot10^{-1}$ | $1.0\cdot10^{0}$ |
| HALLMARK_PI3K_AKT_MTOR_SIGNALING | 0.73 | $9.4\cdot10^{-1}$ | $1.0\cdot10^{0}$ |
| HALLMARK_APICAL_JUNCTION | 0.72 | $9.8\cdot10^{-1}$ | $1.0\cdot10^{0}$ |
| HALLMARK_HEDGEHOG_SIGNALING | 0.63 | $9.7\cdot10^{-1}$ | $1.0\cdot10^{0}$ |
| HALLMARK_ANDROGEN_RESPONSE | 0.58 | $10.0\cdot10^{-1}$ | $1.0\cdot10^{0}$ |
| HALLMARK_UV_RESPONSE_DN | -0.56 | $1.0\cdot10^{0}$ | $1.0\cdot10^{0}$ |
| HALLMARK_NOTCH_SIGNALING | -0.63 | $9.9\cdot10^{-1}$ | $1.0\cdot10^{0}$ |
| HALLMARK_CHOLESTEROL_HOMEOSTASIS | -0.69 | $10.0\cdot10^{-1}$ | $1.0\cdot10^{0}$ |
| HALLMARK_TGF_BETA_SIGNALING | -0.70 | $9.9\cdot10^{-1}$ | $1.0\cdot10^{0}$ |
| HALLMARK_WNT_BETA_CATENIN_SIGNALING | -0.75 | $9.3\cdot10^{-1}$ | $1.0\cdot10^{0}$ |
| HALLMARK_MYC_TARGETS_V1 | -0.76 | $1.0\cdot10^{0}$ | $1.0\cdot10^{0}$ |
| HALLMARK_IL2_STAT5_SIGNALING | -0.80 | $1.0\cdot10^{0}$ | $1.0\cdot10^{0}$ |
| HALLMARK_EPITHELIAL_MESENCHYMAL_TRANSITION | -0.80 | $10.0\cdot10^{-1}$ | $1.0\cdot10^{0}$ |
| HALLMARK_MYC_TARGETS_V2 | -0.81 | $9.0\cdot10^{-1}$ | $1.0\cdot10^{0}$ |
| HALLMARK_COAGULATION | -0.86 | $9.3\cdot10^{-1}$ | $1.0\cdot10^{0}$ |
| HALLMARK_PANCREAS_BETA_CELLS | -0.91 | $6.1\cdot10^{-1}$ | $9.4\cdot10^{-1}$ |
| HALLMARK_APICAL_SURFACE | -0.92 | $6.2\cdot10^{-1}$ | $9.4\cdot10^{-1}$ |
| HALLMARK_DNA_REPAIR | -0.99 | $4.5\cdot10^{-1}$ | $7.3\cdot10^{-1}$ |
| HALLMARK_ESTROGEN_RESPONSE_LATE | -1.00 | $4.4\cdot10^{-1}$ | $7.3\cdot10^{-1}$ |
| HALLMARK_INFLAMMATORY_RESPONSE | -1.01 | $3.8\cdot10^{-1}$ | $6.6\cdot10^{-1}$ |
| HALLMARK_ESTROGEN_RESPONSE_EARLY | -1.03 | $3.6\cdot10^{-1}$ | $6.5\cdot10^{-1}$ |
| HALLMARK_APOPTOSIS | -1.06 | $2.4\cdot10^{-1}$ | $4.5\cdot10^{-1}$ |
| HALLMARK_COMPLEMENT | -1.06 | $2.3\cdot10^{-1}$ | $4.5\cdot10^{-1}$ |
| HALLMARK_E2F_TARGETS | -1.08 | $2.0\cdot10^{-1}$ | $4.1\cdot10^{-1}$ |
| HALLMARK_MYOGENESIS | -1.09 | $1.7\cdot10^{-1}$ | $3.5\cdot10^{-1}$ |
| HALLMARK_GLYCOLYSIS | -1.09 | $1.6\cdot10^{-1}$ | $3.4\cdot10^{-1}$ |
| HALLMARK_ADIPOGENESIS | -1.10 | $1.4\cdot10^{-1}$ | $3.2\cdot10^{-1}$ |
| HALLMARK_HEME_METABOLISM | -1.12 | $1.3\cdot10^{-1}$ | $3.0\cdot10^{-1}$ |
| HALLMARK_UV_RESPONSE_UP | -1.16 | $9.9\cdot10^{-2}$ | $2.5\cdot10^{-1}$ |
| HALLMARK_ALLOGRAFT_REJECTION | -1.16 | $6.5\cdot10^{-2}$ | $2.0\cdot10^{-1}$ |
| HALLMARK_SPERMATOGENESIS | -1.16 | $9.7\cdot10^{-2}$ | $2.5\cdot10^{-1}$ |
| HALLMARK_MTORC1_SIGNALING | -1.17 | $4.7\cdot10^{-2}$ | $1.6\cdot10^{-1}$ |
| HALLMARK_PEROXISOME | -1.18 | $9.5\cdot10^{-2}$ | $2.5\cdot10^{-1}$ |
| HALLMARK_PROTEIN_SECRETION | -1.21 | $6.8\cdot10^{-2}$ | $2.0\cdot10^{-1}$ |
| HALLMARK_KRAS_SIGNALING_DN | -1.22 | $3.3\cdot10^{-2}$ | $1.3\cdot10^{-1}$ |
| HALLMARK_FATTY_ACID_METABOLISM | -1.24 | $3.7\cdot10^{-2}$ | $1.3\cdot10^{-1}$ |
| HALLMARK_MITOTIC_SPINDLE | -1.26 | $4.2\cdot10^{-3}$ | $2.1\cdot10^{-2}$ |
| HALLMARK_UNFOLDED_PROTEIN_RESPONSE | -1.36 | $1.2\cdot10^{-2}$ | $5.1\cdot10^{-2}$ |
| HALLMARK_HYPOXIA | -1.39 | $3.9\cdot10^{-3}$ | $2.1\cdot10^{-2}$ |
| HALLMARK_G2M_CHECKPOINT | -1.41 | $4.1\cdot10^{-3}$ | $2.1\cdot10^{-2}$ |
| HALLMARK_P53_PATHWAY | -1.42 | $3.9\cdot10^{-3}$ | $2.1\cdot10^{-2}$ |
| HALLMARK_TNFA_SIGNALING_VIA_NFKB | -1.50 | $4.1\cdot10^{-3}$ | $2.1\cdot10^{-2}$ |
| HALLMARK_BILE_ACID_METABOLISM | -1.52 | $1.3\cdot10^{-3}$ | $2.1\cdot10^{-2}$ |
| HALLMARK_OXIDATIVE_PHOSPHORYLATION | -1.54 | $3.6\cdot10^{-3}$ | $2.1\cdot10^{-2}$ |
| HALLMARK_REACTIVE_OXYGEN_SPECIES_PATHWAY | -1.63 | $4.9\cdot10^{-3}$ | $2.2\cdot10^{-2}$ |
| HALLMARK_INTERFERON_GAMMA_RESPONSE | -1.73 | $3.7\cdot10^{-3}$ | $2.1\cdot10^{-2}$ |
| HALLMARK_XENOBIOTIC_METABOLISM | -2.02 | $3.5\cdot10^{-3}$ | $2.1\cdot10^{-2}$ |
| HALLMARK_INTERFERON_ALPHA_RESPONSE | -2.08 | $1.1\cdot10^{-3}$ | $2.1\cdot10^{-2}$ |
